# Backstroke rowing as an adaptation for low-Reynolds-number flight in tiny whiteflies (*Bemisia tabaci*)

**DOI:** 10.64898/2026.11.15.766885

**Authors:** Amir Sarig, Lior Gurka, Maor Shemesh, Gal Ribak

## Abstract

The force resisting gravity during flight is typically aerodynamic-lift generated by the wings. However, in the smallest insects the low Reynolds number associated with a wing size ≤ 1mm reduces the efficiency of using lift for flight. We examined how miniature whiteflies (*Bemisia tabaci*) modulate their flapping kinematics to control flight in the vertical plane. Free-flying whiteflies were recorded using high-speed video cameras, and their 3D wing flapping kinematics were analyzed as a function of flight speed and climb angle. Whiteflies displayed the crescent-shaped wingtip trajectory typical to the drag-utilizing propulsion previously described in other miniature insects. To fly faster, they increased the wing sweep amplitude during the upstroke, resulting in increased wingtip speed relative to the body. To fly at steeper climbing angles the insects oriented their body to direct the upstroke wing motion downwards at a high angle-of-attack. Robotic force measurements confirmed that this downwards rowing in air provides vertical aerodynamic force exceeding body weight at least twofold. To direct the upstroke wing motion into a downwards rowing stroke, the insects used a combination of body pitch and roll that often resulted in the insects flying upside-down. This atypical flight posture emphasizes the adjustments required from tiny insects to convert a flight apparatus that was originally intended for utilizing lift into a drag-utilizing propulsion system, enabling flight at low Reynolds numbers.

**Significance statement:** As wings become smaller and flight speed lower, the influence of air viscosity on the flow around the wings becomes proportionally higher, imposing constraints on lift production. To compensate, miniature (wing length ≤1 mm) insects flap their wings in a curved wingtip trajectory that utilizes the drag of the wings for propulsion. We demonstrate that free-flying whiteflies employ a "backstroke rowing" posture, as if swimming on their back through the air during steep ascending flight. Measurements on robotic wings indicate that this backstroke enables vertical force production exceeding body weight 2-4fold. This unique behavioral adaptation enables miniature insects to convert the lift-based propulsion mechanism used by larger insects into a drag-based mechanism that permits flight at the sub-millimeter scale.

## Introduction

Upward flight (climb) is fundamental for take-off (Marden, 1987), reaching high-altitude air currents (Blackmer and Byrne, 1993), maneuvering (Ortega-Jiménez & Dudley, 2018), predator-prey interactions (Le Roy et al., 2022) and even landing in some cases (KleinHeerenbrink et al., 2022). Insects can transition from hovering to climbing by increasing the lift forces they produce to exceed their body weight. In doing so, the wings of an upward moving insect encounter vertical inflow of air onto their stroke plane. Actuator disc (Froude’s momentum) theory shows that this vertical inflow attenuates the efficiency of vertical momentum transfer requiring a higher power input with increasing climb rate (vertical speed) in helicopters (Bramwell et al., 2001, pp 33-76) and insects (Liu and Sun, 2024). Consequently, climbing requires a higher power investment compared to horizontal flight at the same speed (Berg and Biewener, 2008). The added power requirement results in a limitation on climb rate (Parslew, 2015, Johnson, 1980; Liu and Sun, 2024) that can be expressed as a maximal climb angle for a given flight speed or a maximal flight speed for a given climb angle.

Flying animals typically increase their aerodynamic power output by flapping their wings faster through air. The faster wing speed can be achieved by increasing wingbeat amplitude (e.g., Altshuler et al., 2005; Davidovich and Ribak, 2018; Meng et al., 2017) and/or wingbeat frequency (e.g., Davidovich and Ribak, 2016; Krishnan et al., 2022). However, an increase in wing speed alone might be insufficient for the case of climbing flight since the vertical inflow leads to a decrease in the effective angle-of-attack (AoA), thereby lowering the aerodynamic lift produced (Tuan et al., 2020). Consequently, adjustments for climbing flight should also include modulation of the AoA via tilting the stroke plane and/or altering the wing incidence and deviation angles (Dudley, 2000; Ellington, 1984a). While the mechanics of ascending flight has been studied before in flying vehicles (Li et al., 2023), birds (Krishnan et al., 2022), bats (MacAyeal et al., 2011) and insects (Meng et al., 2017), the mechanism enabling such flight in very small insects remains obscure.

Miniature insects (wing length ≤1mm) beat their wings at Reynolds numbers (Re) below 40 (Cheng and Sun, 2021; Farisenkov et al., 2021a; Sane, 2016; Sun, 2023). At these low Re numbers (Re = *c̅U̅_rg_*/*ν*, where *c̅* is the mean wing chord, *U̅_rg_* is the mean speed at the wing’s radius of gyration and *ν* is kinematic viscosity of air), air viscosity imposes limitations on lift production.

Concurrently, the drag coefficient of the wing increases at lower Re, leading to a drop in the lift-to-drag ratio of the airfoil. Low lift-to-drag ratios typically result in low energetic efficiency for flight (Horridge, 1956; Vogel, 1988; Jones et al., 2015), presumably leading to higher power requirement by the insect. Therefore, the power limitation on climb rate may be more pronounced in miniature insects compared to larger ones (e.g. insects of similar size or larger than fruit flies, with wing length >3mm and flight at Re>100). Nevertheless, miniature insects have adapted for flight at the low Re flow regime. Their flapping kinematics differ from that of larger insects in utilizing the high drag on the wings for thrust production (Farisenkov et al., 2021b; Sun, 2023). The wingtip trajectory of miniature insects is curved and asymmetric between the upstroke and downstroke, (Cheng and Sun, 2016; Cheng and Sun, 2018; Williams et al., 2025) forming a crescent-shape. Concurrently, the AoA changes during the flapping cycle to include an impulsive drag-based rowing phase at the beginning of the upstroke when the wing is at high AoA. The upstroke ends in a phase of contralateral wing-wing interaction at the dorsal stroke reversal (’clap- and-fling’, Weis-Fogh, 1973). The fling, terminates with the wing at a large AoA as the wings start the downstroke. Thus, the fling-downstroke transition forms another, yet smaller, rowing phase at the beginning of the downstroke (Cheng and Sun, 2018; Lyu et al., 2019). During hovering, the insect’s longitudinal axis is almost vertical and the two rowing phases at stroke reversals are directed downwards, providing vertical upthrust from drag. Between stroke reversals, the horizontal motion of the wings at mid-stroke may generate some vertical force associated with lift (Sun, 2023). This drag-utilizing flapping differs from the lift-based flapping of larger insects in both kinematics and aerodynamics, raising the question how do miniature insects modulate the vertical force balance required for hovering to perform climbs at different speeds and angles?

To answer this question, we analyzed the climbing flight of sweet potato whiteflies (*Bemisia tabaci,* Hemiptera: Aleyrodidaem, **Fig. 1**). The body and wings of *B. tabaci* are ∼1 mm in length (Ribak et al., 2016). Whiteflies have been reported to be capable of prolonged upwards flight, presumably aimed at reaching high-altitude winds for long-distance dispersal (Blackmer and Byrne, 1993; Byrne, 1999; Wainwright et al., 2017; Wainwright et al., 2020). We hypothesized that whiteflies will modulate their flapping kinematics according to climb rate to account for the additional power needed; but that this modulation capitalizes on drag-based wing rowing rather than wing lift for upthrust. To test this hypothesis, we recorded whiteflies free-flying at different climb angles (Fig 1) with high-speed cameras and analyzed their 3D wing flapping motion as a function climb rate and climb angle. We show that part of the adaptation for climbing flight in these miniature insects involves flying at an up-side down orientation.

**Figure 1:**
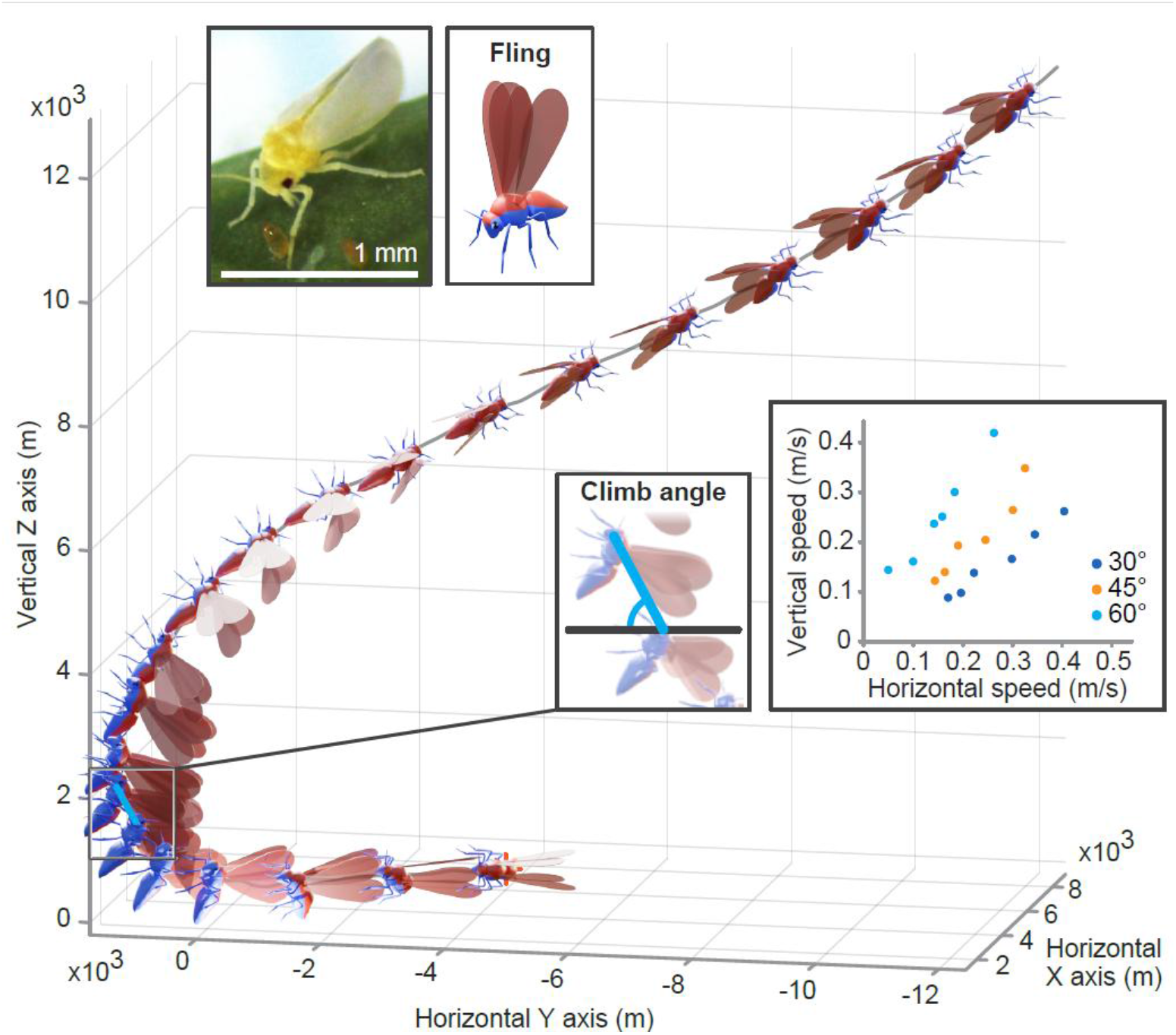
Climbing flight of *Bemisia tabaci*. A) An example of a 3D flight trajectory from one of the movies demonstrating a climb. Animated insect images show the instantaneous position and orientation of the insect during the dorsal stroke reversal (fling) of each flapping cycle. The upper inserts display a picture of a whitefly with scale bar and the colors used in the animation below to distinguish between the dorsal (red) and ventral (blue) surfaces of the insect. The two lower inserts show the definition of climb angle for a single flapping cycle and the horizontal and vertical components of mean flight velocity in the analyzed sequences. Color in the latter denotes the affiliation to three climb-angle categories.

## Results

### Flight kinematics

From the high-speed films of free flying insects (e.g., Supplemental Movie), we selected 18 videos showing the insects climbing at 30°, 45° and 60° climb angles at different flight speed (Fig 1) and subjected these for detailed analysis of the wing and body kinematics. Horizontal flight speed (*U̅_F_*_(*XY*)_) in the selected films ranged from 0.056 m/s for the slowest flight to 0.401 m/s for the fastest one. The climb rate (*U̅_F_*_(*Z*)_) ranged from 0.088 m/s to 0.419 m/s (Fig. 1). This variance in flight velocity was not evident in the mean (± sd) wingtip velocity (*U̅_W_*=1.251±0.124 m/s) or the flapping frequency of the insects that ranged between 167 to 214 Hz and did not correlate with climb rate, climb angle or 3D flight speed (0.02<r<0.09, p>0.9 in all cases). The wing chord-based Re in all the movies was 18±3.5. The advance-ratio increased with flight speed from 0.14 in the slowest flight to 0.41 in the fastest one (Supplemental Information, SI1).

### Modulation of flapping kinematics

The flapping cycle starts with a fling and is immediately followed by the downstroke. The latter ends in the middle of the ventral stroke reversal and the upstroke commences, followed by the clap and then fling (**Fig. 2**). The wingtip path relative to the body was C-shaped in both the upstroke and downstroke. The upstroke wingtip trajectory was consistently posterior to the downstroke one in the body reference frame (BRF), resulting in a crescent-shaped wingtip trajectory for the entire flapping cycle (**Fig. 3**). The wing flapping kinematics of each insect in the BRF were converted to three time-varying Euler angles. The temporal changes in these angles over one flapping cycle are shown in **Figure 4** for the various flight speeds and climb angles. The instantaneous wing sweep angle (**φ**, angular displacement of the wing in the dorsal body plane) has two distinct local minima within the stroke cycle (Fig. 4A). The first minimum is reached at the middle of the downstroke (*t*/*T*=∼0.25) and is moderate (-17.6°±13.39°) compared to the second one (-44.7°±13.95°) which occurs at the end of the upstroke (*t*/*T*=∼0.75). The second minimum (ϕ_min_) became more negative as flight speed (*U̅_F_*) and climb rate increased (r=-0.565, p=0.015 and r=-0.477, p=0.045, respectively, **Fig 5A**), reflecting a more posterior upstroke wingtip trajectory relative to the body in faster flights and climbs. The wingtip speed in the BRF (*u_w_*) averaged over the upstroke duration (0.5< *t*/*T* <0.77) increased as ϕ_min_ decreased (r=-0.603, p=0.008, Fig 5B). Lower ϕ_min_ values were also associated with a larger upstroke inclination in the BRF (*α*, defined in Fig 3, r=-0.8332, p<0.001, Fig. 5C).

**Figure 2:**
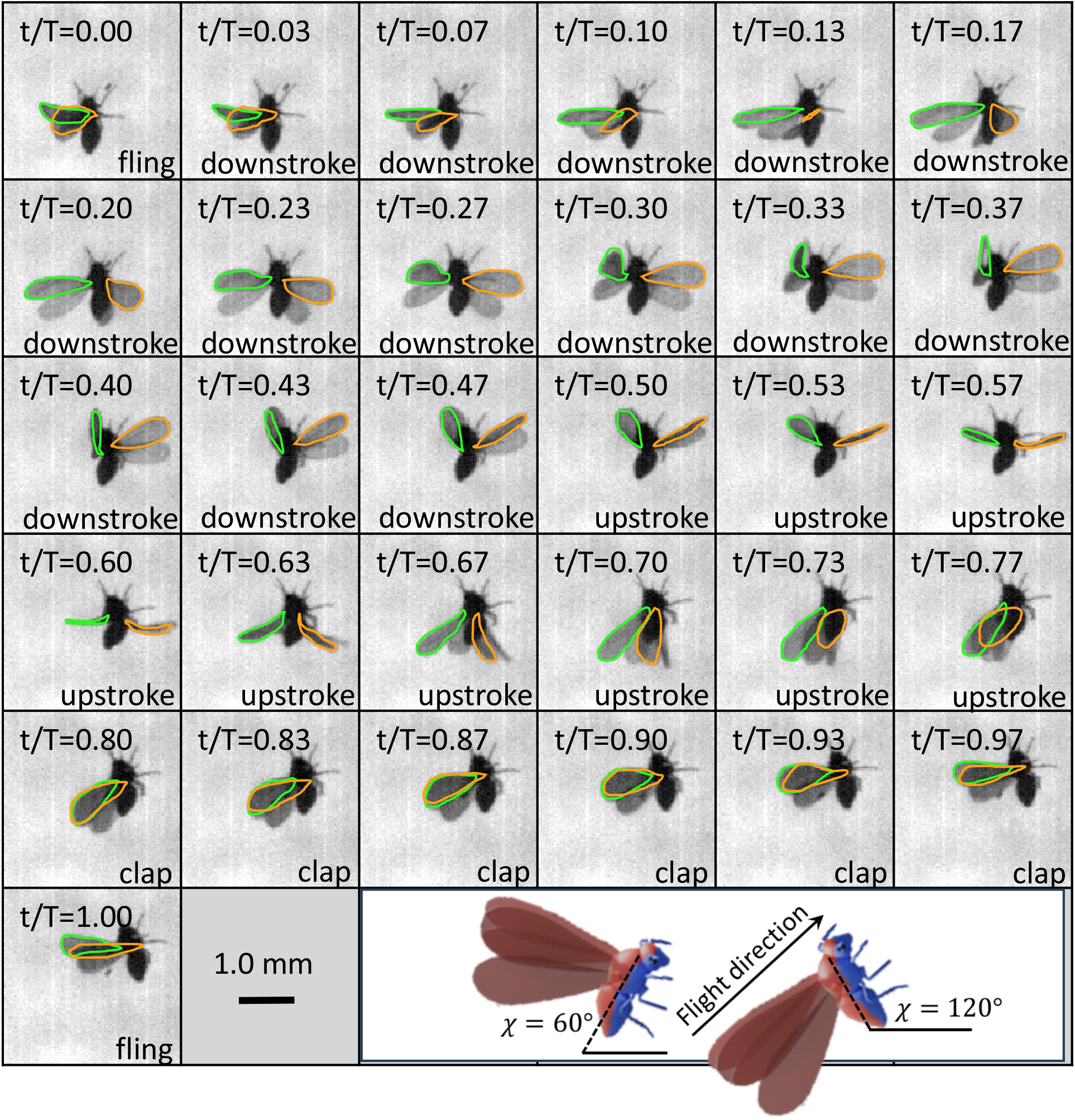
The flapping cycle. High-speed video frames from a single flapping cycle. The time interval between frames is 0.167 ms. The wing contours of the left and right forewings are superimposed on the images. The stroke cycle starts with the fling that marks the beginning of the downstroke. *t/T* is non-dimensional time within the flapping cycle (0≤ *t/T* ≥1). The insert in the lower panel animates the definition of the acute and obtuse body flight angles. The inclination of the longitudinal body axes relative to the horizontal plane is the same in the two examples but the angle is measured with respect to the insect’s ventral (blue) side to distinguish right-side up (*χ* ≤ 90°) and up-side down (*χ* > 90°) body angles.

**Figure 3.**
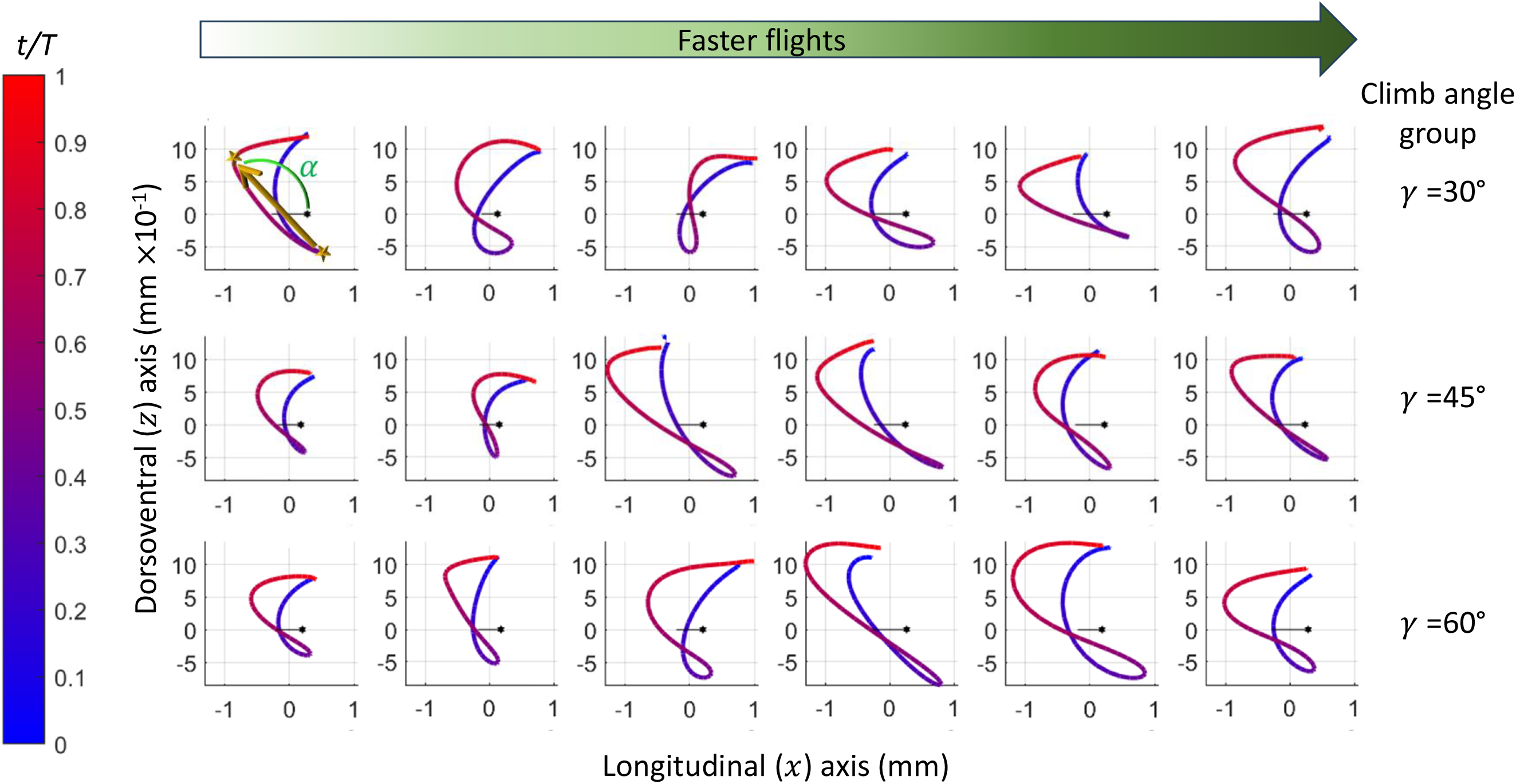
Wingtip trajectory of the analyzed flapping cycles in the body frame of reference. Flapping cycles are arranged according to climb angle (*γ*) category (rows) and 3D flight speed (columns, increasing from left to right). Black horizontal lines and hexagons denote the longitudinal body axis and head positions in each plot. Normalized time within the flapping cycles (*t/T*, starting with the fling) is denoted by the line color according to the left color bar. In the upper left plot, stars indicate the position of the minimum wing-elevation angle and the minimum wing-sweep angle that define the upstroke initiation and termination, respectively. A vector connecting the two positions (***p_upstroke_***, yellow arrow) defines the upstroke inclination (e.g., in eq. 1 and 2). The angle *α* in the sagittal plane between this vector and the longitudinal body axis is shown in green.

**Figure 4:**
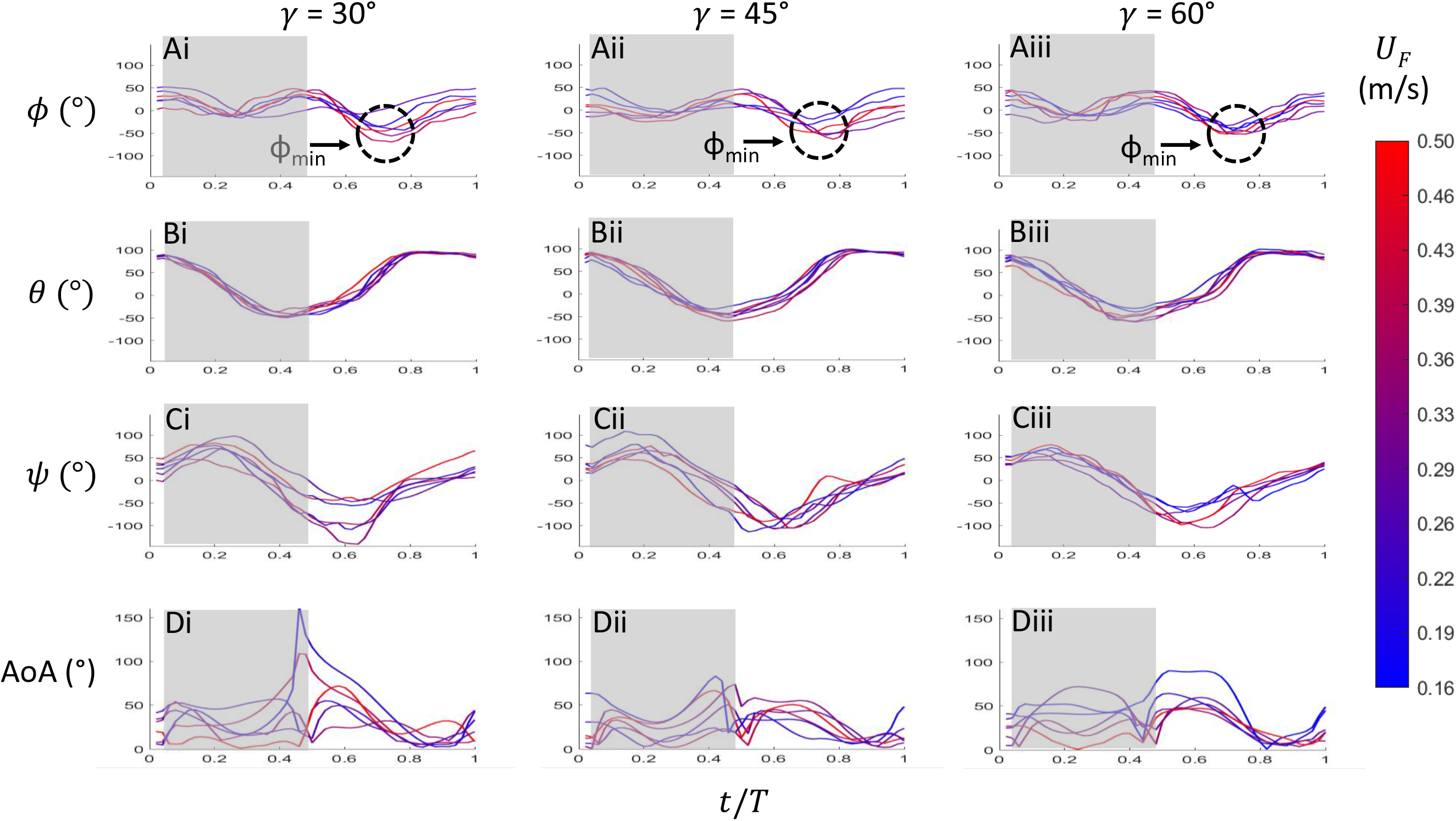
Change in wing flapping angles and angle-of-attack during the flapping cycle. A-C) *φ*, *θ* and *ψ* correspond to sweep, elevation and incidence angles, respectively, measured in the body reference frame. D) AoA corresponds to the wing’s angle-of-attack measured in the inertial world reference frame. Columns i-iii correspond to the three climb angles (*γ*) categories. Each curve represents data from one film with line color denoting flight speed according to the color-bar on the right. t/T, is the non-dimensional time within the flapping cycle (0-1). Grey backgrounds denote the period of the downstroke. Dashed circle in A mark the second sweep minimum ϕ_min_, which correlated with the flight speed and climb rate.

**Figure 5:**
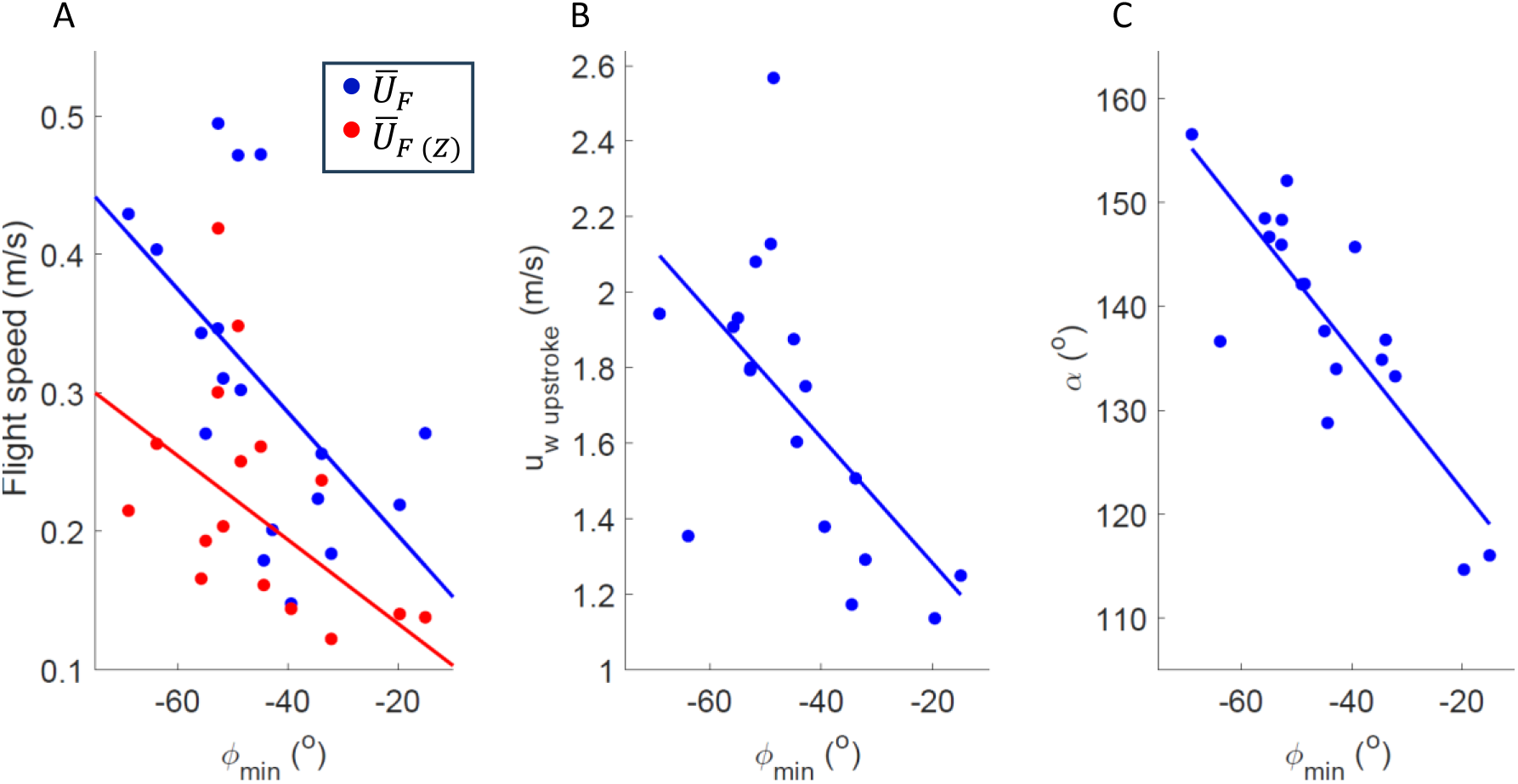
Correlation between wing sweep minimum (*φ_min_*, measured in the body reference frame) and: A) 3D flight speed (*U̅_F_*, blue) and climb rate (*U̅_F_* _(*Z*)_, red) in the world reference frame, B) mean 3D wingtip speed during the upstroke measured in the body reference frame, C) the upstroke inclination angle (*α*) measured in the body reference frame.

In contrast, the pattern of the other Euler angles, wing elevation (*θ*, angular displacement in the sagittal plane, Fig. 4B) and wing incidence (*ψ*, angular displacement of the wing about its length, Fig. 4C), included minimum and maximum values that did not significantly correlate with the climb rate, climb angle or *U̅_F_* in the movies (Supplemental Information, SI1).

The wing’s angle-of-attack (AoA, Fig. 4D) was measured in the inertial world reference frame (WRF). It varied considerably across the analyzed sequences and within climb angle groups. It consistently diminished between the fling and the first sweep angle minimum (0.1<*t*/*T*< 0.3). It then gradually increased, approaching 90° at the ventral stroke reversal (*t*/*T*=0.45). Following wing supination, the AoA remained high as the wings moved during the upstroke up to ϕ_min_ at *t*/*T*=∼0.75, and then reached low values (0°<AoA<20°) as the upstroke transitioned to the clap phase. The clap ended with an abrupt increase in the AoA during the fling. During the early upstroke, (0.5<*t*/*T*<0.7), the AoA was consistently high (>45°) at the 60° climb angle group (Fig. 4Diii). However, we did not find a significant correlation between the mean upstroke AoA and climb rate or climb angle (-0.13<r<0.18, p>0.48 for both cases).

The wingtip speed in the BRF (*u_w_*) was minimal at the ventral stroke reversal but quickly reached its peak value during the upstroke (*t*/*T*=∼0.6), prior to the wing sweep reaching ϕ_min_ (*t*/*T*=∼0.75). Concurrently, the vertical component of wingtip velocity measured in the WRF (*U_W_*_(*Z*)_) had a negative peak at *t*/*T*=∼0.6, indicating that the wing was moving down while at top speed (**Fig. 6**).

**Figure 6:**
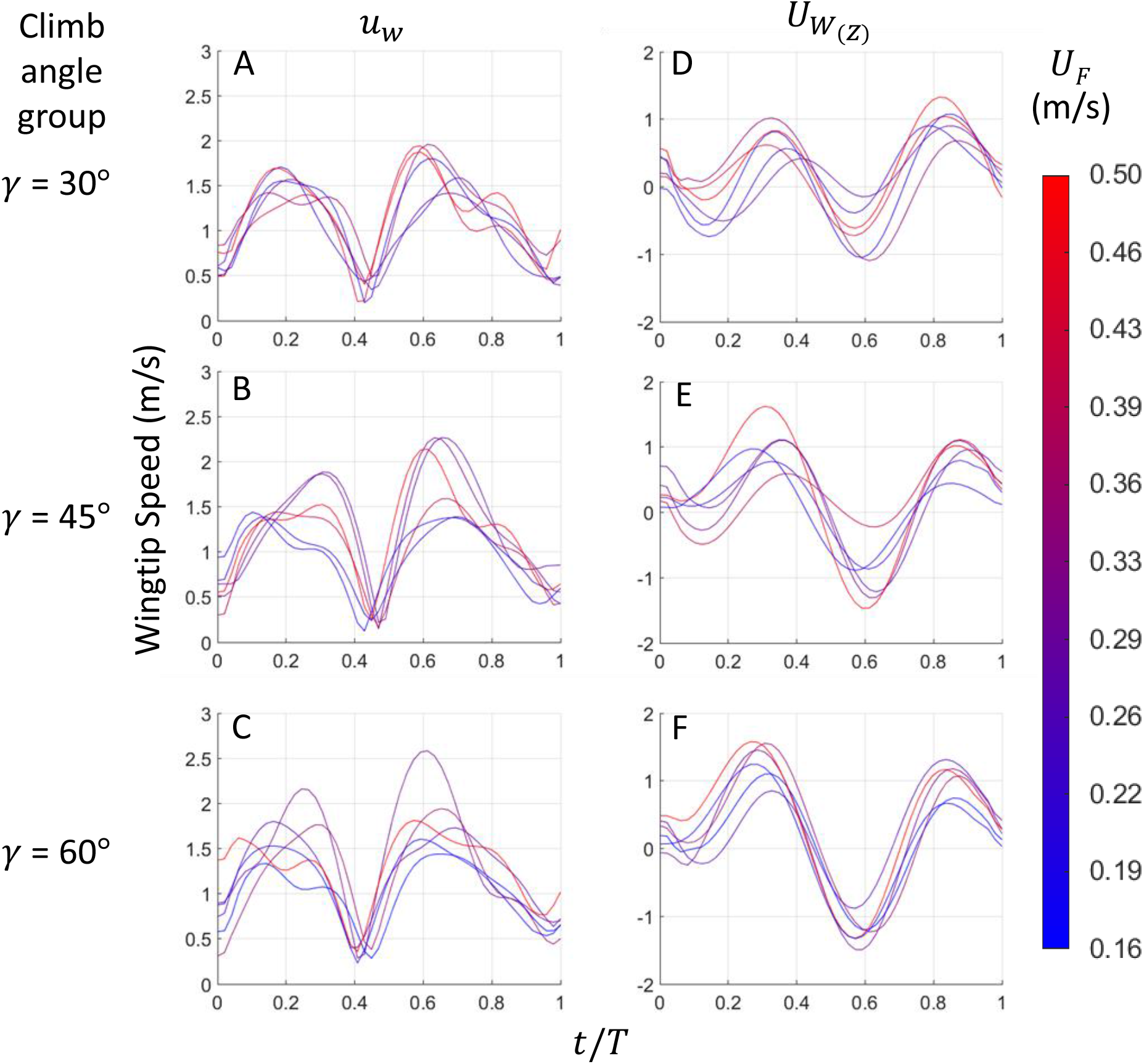
Changes in wing speed during the flapping cycle. Rows correspond to different climb angles. The left and right columns correspond to 3D wingtip speed (*u_w_*) in the body reference frame and the vertical component of wingtip speed (*U_W_*_(*Z*)_) in the world reference frame, respectively. Each curve depicts data from one film with line color denoting the mean 3D flight speed of the insect according to the color-bar on the right. *t/T* is the normalized time within the flapping cycle.

This wing speed minimum was more negative in movies displaying a higher climb angle (r=-0.59, p=0.01). A second, less prominent, downward wing movement (*U_F_*_(*Z*)_<0) was associated with the transition from the fling to the early downstroke (*t*/*T*=∼0.1). It however diminished in steeper climb angles (Fig. 6D-F). A lower minimum *U_F_*_(*Z*)_ at *t*/*T*=∼0.6 was not necessarily accompanied with an increase in peak *u_w_* at *t*/*T*=∼0.6. Rather, the increase in downwards wing speed was mostly due to the body orientation in the WRF as explained next.

### Body orientation and redirecting the flapping motion

The whiteflies in the different movies flew at a broad range of body angles (40°≤ χ ≤152°, mean = 101°± 41°), with obtuse angles being common at all climb angles but dominating in the 60° climb angle group (**Fig 7A**). Although χ increased and its variance across films decreased with increasing climb angle group (Fig. 7A), χ did not statistically differ between the three climb angle groups (Kruskal-Wallis ANOVA H_(2,N=18)_=2.558, p=0.278) due to some of the insects at the lower climb angle groups flying with obtuse body angle as well. At the obtuse body angles, the whiteflies were flying upside down, with the head leading in the flight direction but their ventral surface and legs facing upwards in the WRF. Body pitch showed a weak negative correlation with body roll (r=-0.46, p=0.052, Fig 7B) and body roll showed a weak positive correlation with climb angle (r=0.46, p=0.057), although neither correlation was statistically significant at the 95% confidence level.

**Figure 7:**
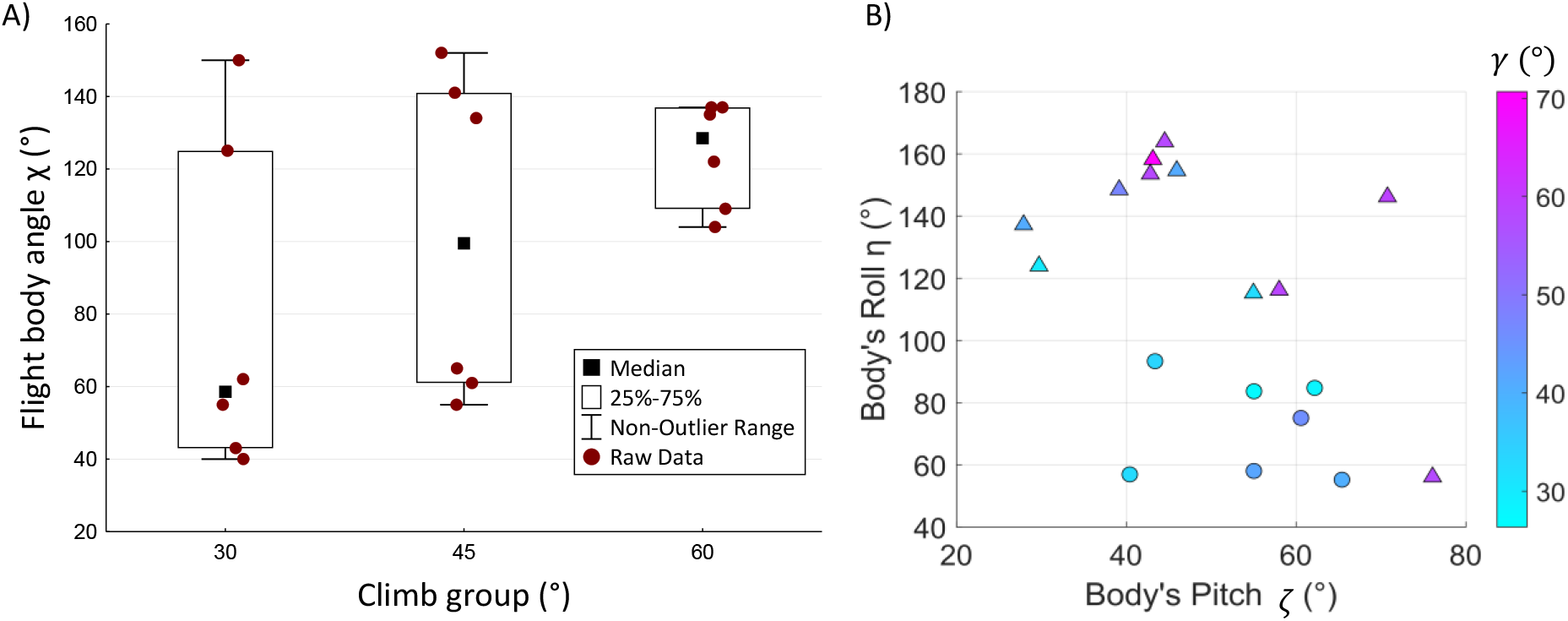
Body orientation. A) The mean body angle for each film (dots) tended to increase while its variance between films decreased with increasing climb angle groups, although the trend was not statistically significant (Kruskal-Wallis ANOVA H_(2,N=18)_=2.558, p=0.278). B) The relationship between mean body pitch (*ξ*) and body roll (η) in the different video sequence. Circles and triangles denote cycles in which the insects are right-side up and up-side down, respectively. Symbol colors denote the climb angle (*γ*) according to the color-bar on the right.

The different body orientations combined with the flapping kinematics resulted in the wings moving downwards (in the WRF) during the upstroke. To verify the prevalence of this phenomenon we filmed and examined the climbing flight of another 15 insects flying at climb angles 40° – 70°. Visual inspection of the footage confirmed that 10 out of the 15 observed insects climbed at *χ*>90° with their wings moving downwards and their tips pointing towards the ground during the upstroke (Supplemental Information, SI2). **Figure 8** exemplifies the kinematics of the downwards rowing in the WRF, in the three fastest flights of each climb angle category. The beginning of the upstroke is denoted by dashed ellipses emphasizing the downwards wing movement at a large AoA. This vertical rowing was affected by body orientation, as evident in

**Figure 8:**
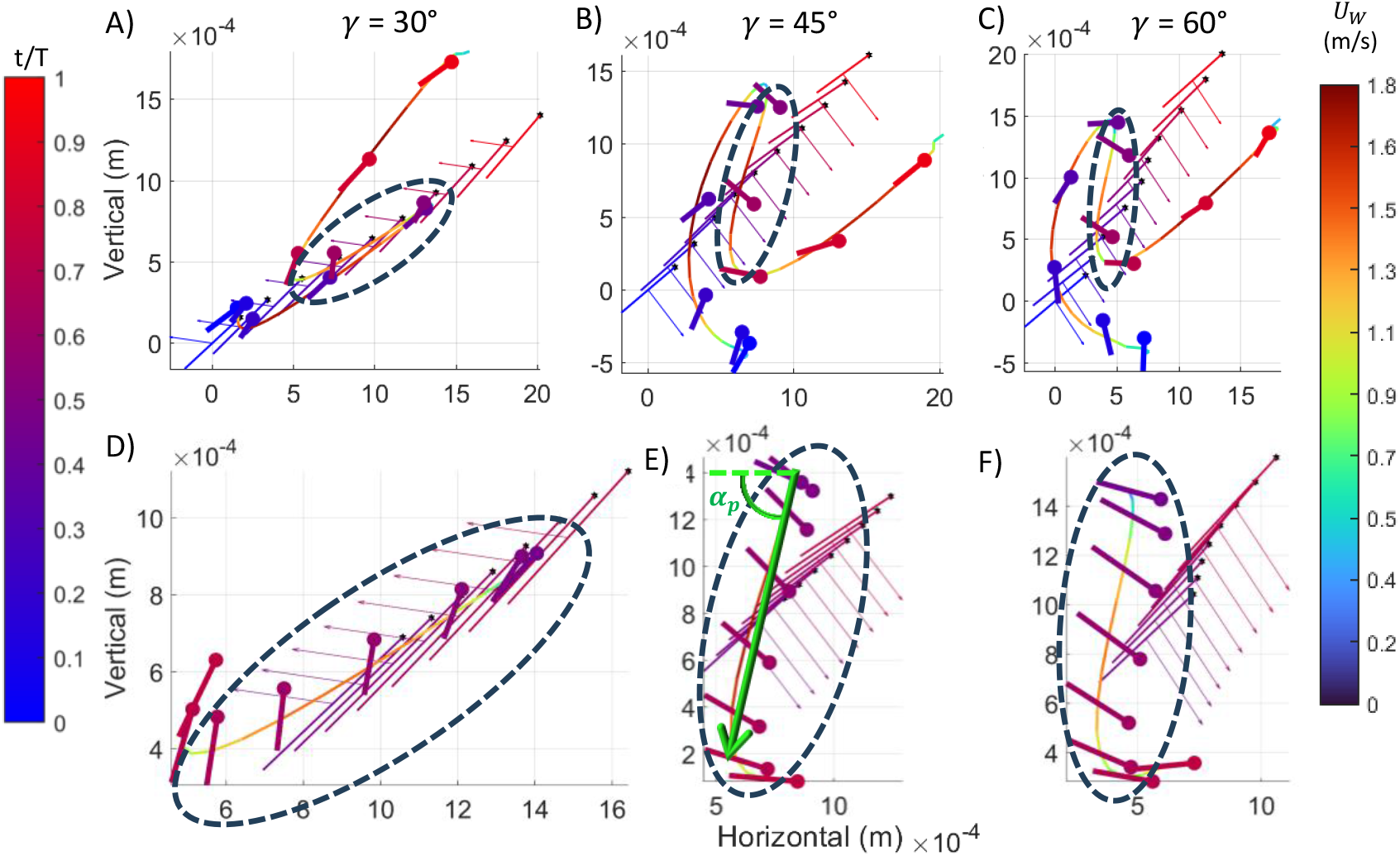
Wingtip trajectory and body orientation in the world reference frame. The plots depict the fastest flight in each climb angle (*γ*) category A) 30°, B) 45°, C) 60°. D-F show the same data as in A-C, respectively, but zoomed in on the upstroke (0.5< *t*/*T* <0.75, dashed ellipses). The horizontal flight direction is to the right (towards the positive side of the horizontal axis). The wingtip trajectory is color-coded according to the 3D wingtip speed in the world reference frame (right color-bar). The body’s longitudinal axis (thin lines) and wing chord (thick lines) are depicted each 0.1 *t*/*T* in A-C and each 0.04*t*/*T* in D-F. They are color-coded according to *t*/*T* (left color-bar), with circles denoting the head and the wing’s leading edge, respectively. Arrows perpendicular to the body’s longitudinal axis point to the dorsal side of the body. Note that in A the whitefly is flying right-side up with its ventral body surface facing the flight direction. In contrast the insects in B and C are flying up-side down. The green arrow in E illustrate the definition of the downwards rowing direction (*α_p_*). Wingtip trajectories of the remaining 15 flapping cycles are displayed in Supplemental Information, SI3.

Fig. 8B and 8C, where the whiteflies are flying upside down; whereas in Fig. 8A the insect is flying right-side up (See also Supplemental Information, SI3). The climb angle in each movie was correlated with the direction of the upstroke wing motion measured in the WRF (***α****_p_*, defined in Fig 8E, r=-0.69, p=0.002). At steeper climb angles, ***α****_p_* approached -90°, i.e., making the wing movement at this phase more vertical and downwards directed.

### Robotic force measurements

We replicated the mean flapping kinematics of each climb angle group in a dynamically scaled robotic wings system to measure the fluid forces associated with the observed flapping kinematics (**Fig. 9**). The force measurements revealed that the upstroke rowing yielded high drag forces that combined with smaller lift had a vertical component that exceeded the estimated body weight of the insects (<40 µg, Ribak et al. 2016) at least twofold in the 30° climb angle and 3-4fold in the 45° and 60° climb angle groups (Fig. 9E).

**Figure 9.**
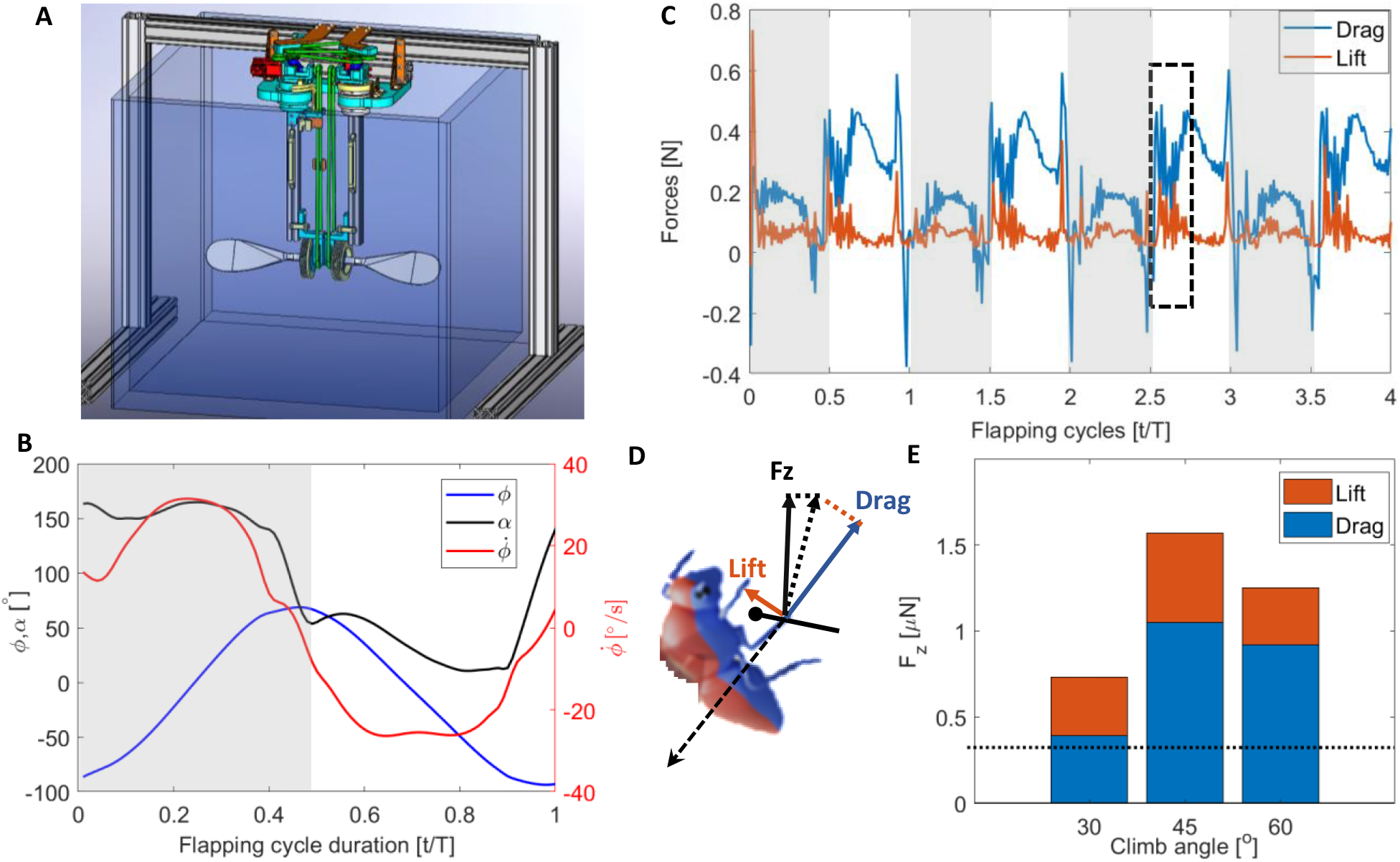
Robotic measurements of aerodynamic force. A) the robotic setup includes two upscaled forewing models actuated at two degrees of freedom through Glycerol. A 3-axes force transducer at the root of the wing-model measured the axial and normal forces on the wing. B) The robot is programed to move the wing at prescribed flapping angle (*φ*) and angle of attack (*α*) extracted from the high-speed movies, while the speed is dynamically scaled to maintain the same Re numbers. The kinematic shown in the figure are the mean from all flights in the 30° climb angle category. C) Each force measurement includes four flapping cycles in sequence. The measured forces are converted to drag (blue) and lift (orange). The upstroke rowing forces are measured from the first half of the third upstroke (2.5≤ *t*/*T* ≤2.75, dashed black rectangle) to allow flow development. Gray and white backgrounds correspond to downstrokes and upstrokes, respectively. D) The instantaneous vertical force (Fz) used to overcome gravity and move the insect upwards at each instant during the early upstroke is comprised of the lift and drag force. E) When averaged over time during the early upstroke (2.5≤ *t*/*T* ≤2.75) and scaled back to the insect dimensions the vertical force from the four wings greatly exceeds the body weight of whiteflies (the larger females weigh ∼40 *μ*g, dotted horizontal line). Fz is comprised mostly of drag (blue bars) with a smaller contribution from lift (red bars). Each stacked column corresponds to the mean kinematics for insect climbing at 30°, 45° and 60°.

## Discussion

To move up, *B. tabaci* direct their upstroke wing motion downwards, to utilize the high wing drag for rowing the body upwards. This vertical rowing was enabled by two main modifications by the insects: In the BRF, to climb at a higher rate the insects moved their wings posteriorly during the upstroke at a larger sweep angle, as evident by a lower *φ_min_*. This adjustment to the wingtip trajectory resulted in both increased wingtip speed relative to the body during the upstroke and pointed the upstroke direction posteriorly at a larger *α* angle (Fig. 5c). The second modification was the orientation of the body in the WRF with respect to gravity and flight direction. Namely, the insects oriented their body’s pitch and roll, often displaying obtuse body angles that converted the upstroke trajectory in the BRF to a vertical rowing motion in the WRF. These obtuse angles often resulted in the insects flying upside-down; i.e., with their dorsal side facing the ground.

The redirecting of the upstroke direction (***α****_p_*) from the insect’s point of view can be described as:

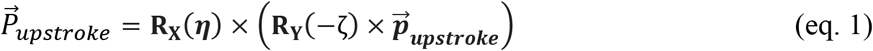

where 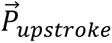 is the upstroke trajectory vector rotated to the WRF, **R_X_** and **R_Y_** are rotation matrices using the body roll and body pitch angles to rotate 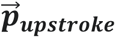 about the X and Y axes respectively. 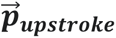 is the upstroke trajectory vector in the BRF (**Fig. 3**). Since ***α****_p_* is the direction of 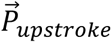 in the vertical plane, it can be measured as the angle between its Z and XY components:

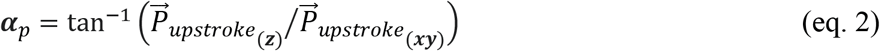

Comparing ***α****_p_* measured directly from the movies (e.g., Fig 8E) with ***α****_p_* calculated using eqs. 1 & 2 showed that 93% of the variation between movies in the observed ***α****_p_* could be explained by body pitch, body roll and the angle *α* describing the direction of 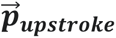 in the sagittal (BRF) plane (Supplemental Information, SI4). Since *φ_min_* correlated with both *α* and *u_w_*, its change contributed to regulating the upstroke rowing phase in the BRF while the body pitch and roll angles reoriented this rowing motion to increase its vertical component in the WRF.

The wingbeat kinematics of *B. tabaci* at different climb angles and climb rates underlines the distinction between miniature insects and larger ones. In insects the size of fruit flies and larger (>3-4 mm), lift is the predominant force for keeping the insect aloft. The flapping frequency, flapping amplitude and AoA are the main parameters that determine this lift (Dickinson et al., 1998). Inclination of the stroke plane in the WRF (typically by adjusting body pitch) then redirects this lift force, which is roughly perpendicular to the stroke plane. In contrast, whiteflies adjusted their climb rate by directing the upstroke rowing motion more vertically in the WRF (Figs. 6, 8, Supplemental Information, SI4) to utilize the drag on the wing for propelling the body upwards. This finding substantiates recent reports on miniature insects using wing drag for supporting their body weight in the air while hovering (Cheng and Sun, 2018; Farisenkov et al., 2021b; Sun, 2023) and extends it to the case of climbing flight.

When the body’s longitudinal axis is vertical (*ζ*=90°) the upstroke motion is downwards and backwards in the WRF leading to a drag-based thrust upwards and forward (Fig. 10). Any lift generated during the upstroke would further contribute upthrust but also a horizontal force component that opposes the horizontal drag-based thrust (Fig 10). To climb at a steeper angle an obtuse *χ* can further tilt the upstroke vertically but now the insect is up-side down, with its ventral surface facing upwards and the tip of the abdomen leading (e.g., *χ*=120° in Fig. 10). Wing lift would shift the upstroke thrust towards the head direction. Even more obtuse *χ* would lead to the insect remaining upside down but now the horizontal thrust direction imply that the insect is moving head first (see *χ*=150° in Fig. 10) with drag providing mostly upthrust and lift providing horizontal thrust. This obtuse body angle is equivalent to the insect flying at a small pitch angle (*ζ*=30°) and a large body roll (*η*=180°), explaining the weak negative relationship observed between *ζ* and *η* (Fig. 6B). For a horizontally-oriented insect (*ζ*=0°), flying upside-down (*η*=180°) is the only way to utilise the upstroke drag for both forward thrust and weight support (see *χ*=180° in Fig. 10). The observed variation in *α* between 114°-155° (Fig 5C), provides further adjustment of ***α****_p_* by up to ∼40° (green arrows in Fig. 10). The increase in *α* is coupled with an increase in *u_w_* during the upstroke through their mutual dependency on *φ_min_* (Fig 5B-C).

**Figure 10.**
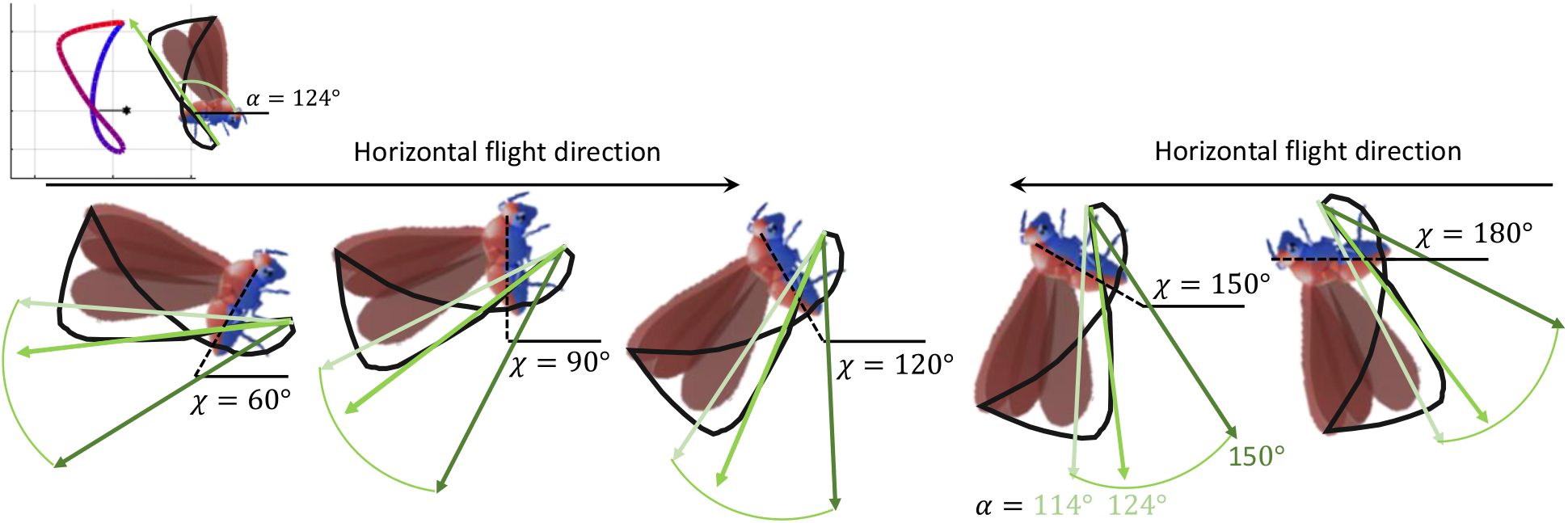
Effect of body angle (*χ*) and *α* values on upstroke direction. Upstroke direction is shown in green arrows for different *α* values.

An obtuse body angle is therefore effective for climbing with the head leading but results in the insect flying up-side down. This was the case in 11 out of the 18 movies analyzed for flapping kinematics and in another 10 out of 15 movies with climb angles *γ*>40° that were inspected visually. Hence, a climb with an up-side down body orientation seems to be more frequent than climbing right-side up, although evidently both are possible (Fig 8).

During the upstroke rowing, *U_W_*_(*z*)_ started from ∼0 ms^-1^ at the ventral stroke reversal (*t/T* =0.4) and reached a peak negative value at *t/T*=∼0.6 (Fig. 5). The acceleration of the wings (in the WRF) during this period ranged between 558 and 2308 m/s^2^. The acceleration vector is roughly parallel to wing velocity, contributing acceleration reaction (wing inertia + added mass) to propel the body in the same direction as wing drag. Wing acceleration has been suggested as the major aerodynamic drag enhancer during the upstroke rowing phase of miniature insects (Sun, 2023). In the present study, the vertical (downwards, i.e. negative) component of wing acceleration during the early upstroke was also found to be correlated with the climb angle (r=-0.557, p=0.016), supporting its role in augmenting upthrust for the climb.

A consequence of *B. tabaci* using wing drag to climb is that the climb speed affects the wings aerodynamics differently than in larger insects and helicopters. In the latter, vertical inflow of air interferes with the induced velocity and downwards momentum transfer of the horizontally moving wings by diminishing the AoA. The resulting reduction in lift can be compensated by faster horizontal wing motions or actively increasing the wing incidence angle. In the drag-based vertical climb of *B. tabaci*, the wings move downwards at a large AoA. This rowing motion is thus less sensitive to changes in AoA compared to lift generating airfoils that rely on attached flows. In *B. tabaci* the geometrical AoA of the wing during the upstroke did not contribute to explaining the climb angle or climb rate. However, we noted that the AoA was always high (>45°) during the rowing phase in the sequences showing the insects flying at the higher climb angle groups compared to the lowest one (Fig. 4D).

The downward rowing of *B. tabaci* is sensitive however to the climb rate since the upwards speed of the insect decreases the downwards speed of the wing during the upstroke in the WRF. This effect is evident by the diminishing first minimum of U_W(Z)_ early in the downstroke (t/T =0.1) with increasing climb angle (Fig. 6). consequently, the fling-associated rowing cannot contribute much for high climb rate. In contrast, the upstroke rowing had a faster *U_W_*_(*Z*)_ at *t*/*T*=0.6 at larger climb angles (Fig. 5). Nevertheless, the effect of upwards body speed on downwards rowing can be seen in Fig. 5 by noting that the *U_W_*_(*Z*)_ during the upstroke did not exceed 1.5 ms^-1^. At the same time *u_w_* reached ∼2 ms^-1^ while the vertical flight speed was < 0.42 ms^-1^.

The whiteflies not only mitigated the reduction in vertical wing speed from upward body motion but they also increased *U_W_*_(*Z*)_ at steeper climbs by increasing *u_w_* and *α* and redirecting ***α****_p_* to have a larger vertical wing speed component during the upstroke. Moving the wing faster relative to air can be done by increasing the flapping frequency, the flapping amplitude, or both. Our whiteflies did not increase their flapping frequency at faster climb rates, climb angles or flight speed. The increase in *u_w_* was instead gained by increasing the wing sweep amplitude during the upstroke allowing to increase the speed of the wing during the rowing phase specifically. In larger insects, maximal vertical flight speed is predominantly determined by the power needed to overcome the drag of the faster moving body and the drag on the faster moving wings. In miniature insects such as *B. tabaci* the upper limit to vertical flight speed is in addition dependent on the decrease in vertical wing rowing speed with increasing climb rates.

The observed flapping kinematics in climbing *B. tabaci* corroborates previous reports on a highly curved wingtip trajectory in these insects (Williams et al., 2025). A similar, crescent-shaped, wingtip trajectory with a clap-and-fling mechanism at the dorsal stroke reversal and a pronounced rowing phase following the ventral stroke reversal has been reported in other miniature insects from different orders such as thrips (Thysanoptera), miniature flies (Diptera) and parasitic wasps (Hymenoptera) (Sun, 2023). The similarity of these kinematic features across miniature insects with low phylogenetic relatedness likely reflect convergent evolution in the adaptation for flapping flight aerodynamics at the <1 mm scale.

Miniature insects would appear to utilize wing drag to propel their body, using a flight apparatus that originally evolved for generating lift. Consequently, the crescent-shaped wingtip trajectory and the use of body orientation to redirect upstroke drag results in flight at extremely variable body (χ = 58°-144°) and roll angles (0-158°) that would be considered abnormal in larger insects, emphasizing the adjustments required for flight in the low Reynolds number range.

## Material and methods

### Insects

Whiteflies (*B. tabaci*, **Fig. 1**) were collected using an aspirator from a population kept in a greenhouse at Tel Aviv University inside fine-mesh cages containing *Sonchus oleraceus* plants. The insects were transferred to glass vials and taken to the laboratory to record their flight on the same day of collection.

### High-speed filming setup

Three high-speed cameras (Fastcam SA3, Photron Inc.) equipped with 85 mm lenses (Nikkor, Nikon) recorded the insects free-flying in stagnant air at 6,000 frames per second (Supplemental Movie). The cameras were spatially calibrated using the MATLAB script DLTcal5 (Hedrick, 2008) by moving an insect mounting pin with a micromanipulator (Thorlabs) to known positions in 3D space. High-intensity near-infrared (850 nm) LED floodlights (Elimec LTD) supplied back-light illumination to the cameras at a wavelength above the detection threshold of most insects (∼700 nm, Döring & Chittka, 2007). We did not observe any directional bias during the flight of the tested insects towards or away from the infrared light source. A glass vial housing the insects with a small opening at its tip enabled them to exit the vial individually. The insects voluntarily began wing-flapping, usually while still inside the glass vial, and exited it through the opening. The opening of the vial was positioned 2.5 cm (∼30 body lengths) below the cameras’ fields of view to ensure that any transient behaviors associated with flight initiation, including the take-off jump (Ribak et al., 2016), were not recorded in our high-speed videos. In total we filmed 524 wing flapping cycles from 58 different individuals.

### Reconstruction of kinematics

In the Cartesian coordinate system of the spatially-calibrated cameras the +Z-axis was parallel to gravity and pointing up while X and Y were horizontal (Fig. 1). We found the instantaneous flight velocity (*U_F_*) of the insects by digitizing the position of the head in each movie frame and applying a numerical time derivative (Rayner and Aldridge, 1985) on the 3D position data. We then calculated the mean flight velocity (*U̅_F_*) for each flapping cycle, and from the 3D components of the mean velocity vector (*U̅_F_*_(*X*)_, *U̅_F_*_(*Y*)_, *U̅_F_*_(*Z*)_) we found the climb angle (*γ*, Fig. 1) for that cycle:

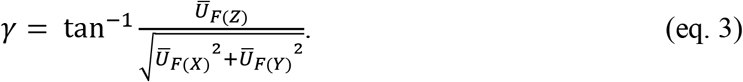

where overbars denote cycle-averaged means. *U̅_F_*_(*Z*)_ represents the climb rate of the insect. We then singled out the flapping cycles with climb angles 30°, 45° and 60° (±7°, Fig. 1) and sorted *U̅_F_* within each angle category to select six flight speeds per climb angle category that represent the observed speed range (see inset in Fig. 1). We gave preference to flight sections that had the lowest accelerations and least deviations from a straight flight trajectory. The resulting 18 flapping cycles, from 18 different individuals, were digitized frame-by-frame to extract the wing-flapping kinematics throughout the flapping cycle.

To overlap the data from different flights we rotated the coordinate system about the Z axis such that the +X-axis coincided with the horizontal component of flight direction and this rotated Cartesian coordinate system (X,Y,Z) is referred to hereafter as the world reference frame (WRF). We also used a coordinate system fixed on the body of the insect (x,y,z) with the origin fixed at the wing hinge. This latter coordinate system is used to describe wing motions relative to the body and referred to as the body reference frame (BRF). The head and thorax define the direction of the longitudinal body axis (with +x pointing towards the head). The abdomen was not included in the longitudinal body axis measurement since it moves vigorously relative to the thorax in each flapping cycle (Williams et al. 2025). The two wing hinges defined the transverse axis, with +y pointing to the left of the animal.

To extract wings and body kinematics we used a previously published MATLAB script (Fontaine et al., 2009; Fry et al., 2003; Fry et al., 2005). The body position and orientation are represented in each movie frame by four points: the tip of the head; the connection between the thorax and abdomen; and the two wing hinges. The position and orientation of the left- and right-wing is then determined in each movie frame using a manually-fitted 2D computer model of the contour of *B. tabaci* forewings. The hindwings of *B. tabaci* resemble the forewings in shape and size. They trail the motion of the forewings (Supplemental Information, SI5) enabling us to focus on describing the motion of the forewings as a general representation of the kinematics of both wing pairs.

### Data analysis

The time series of wing and body positions (WRF coordinates) were smoothed in MATLAB using a Butterworth filter with a cut-off frequency of 600 Hz. From the smoothed position data, we calculated numerical time derivatives (Rayner and Aldridge, 1985) to get instantaneous wingtip velocities and accelerations. We distinguish herein between ‘wingtip velocity’ (*U_W_*) measured in the WRF and ‘wingtip velocity relative to the body’ (*u_w_*) measured in the BRF. The advance-ratio (flight speed/wingtip speed, Ellington, 1984a) for each analyzed flapping cycle was defined as *U̅_F_* divided by *U̅_W_*.

To compare and overlay the wing kinematics data in plots, we: 1) averaged the kinematics of the left and right wings in each flapping cycle, 2) normalized time (*t*) by flapping cycle duration (*T*, measured from the number of movie frames), thus obtaining a non-dimensional timescale within the flapping cycle (0≤ *t*/*T* ≤1), 3) interpolated the data to 50 equally-spaced time steps (step width=0.02 *t/T*) using the cubic-spline function in MATLAB. In our definition, *t*/*T*=0 is the end of the wing clap and beginning of the fling.

The body angle during flight (χ, Ellington, 1984b) was defined in the WRF as the angle between thorax longitudinal axis and the horizontal plane. However, in climbing *B. tabaci* the insects often flew with their dorsal side facing down (Fig. 1). Therefore, we measured χ from the insect’s ventral side to distinguish between acute (χ< 90°) and obtuse (χ>90°) body angles when the insects were flying right-side up and up-side down, respectively (Fig. 2). We also extracted the roll, pitch and yaw angles of the body from the vector cosines matrix of the body axes in the WRF. Since yaw was negligible in the movies, body roll (η) gave the angle between the body’s dorsoventral axis and the vertical (XZ) plane and body pitch (*ζ*) equals χ for insects with zero body roll (right-side up orientation).

To evaluate how the insects modulate their flapping kinematics with climb rate we extracted the forewings’ time-varying Euler angles in the BRF. Supplemental Information, SI6 describes the equations and steps necessary to extract these angles and to rotate the data from the WRF to the BRF. Briefly, we used the three axes of the body in each movie frame to rotate the wings data about their hinges from the WRF onto the BRF. We then used the rotated wing data to find three time-varying wing-flapping angles (ϕ, θ *ψ*) where θ is the wing elevation angle defined as the angle between the wing length (line connecting the wing hinge and tip) and the projection of this line in the insect’s horizontal plane. ϕ is the wing sweep angle defined as the angle between the same wing length and the insect’s transverse axis, after rotating the wing by θ around the insect’s longitudinal axis to the frontal (xy) plane. The incidence angle (*ψ*) is the angle between the wing chord at 0.7 of the wing length and the insect’s xy plane after rotating the wing by θ and ϕ. The angle-of-attack (AoA) of the wing is measured in the WRF as the angle between the wing’s chord and the wing’s velocity vector (*U_W_*). A mean upstroke inclination angle (*α*) was defined in the BRF as the angle between a 3D vector (***p_upstroke_***) connecting the wingtip positions at upstroke initiation and termination and the xy plane (**Fig. 3**). Means are presented ± sd where sample size in all the analyses was n=18 insects.

### Robotic measurements

The contribution of the upstroke motion to climbing and weight support was investigated by replicating the flapping motion on a pair of 290-fold upscaled *B. tabaci* wing models moved through glycerol after dynamically scaling the motion to have the same Reynolds number as the insects. The wing models (Supplemental Information, SI7) were prepared in Solidworks based on the forewing planform of *B. tabaci* (Ribak et al. 2016) and 3D-printed. They were then actuated using a custom-built robotic apparatus in a 1×1 m large and 0.8 m deep glycerol tank. The robotic apparatus had two degrees of freedom allowing to flap the wings horizontally at a prescribed angular velocity while controlling the wings AoA at each instant. The base of one of the robotic arms had a three-axes force transducer (K3D60, Me system) to measure the forces generated by the moving wing model. A second set of 3D-printed models had the same density, mass and position of the center of mass as the wing models, but these models had a cylindrical shape. The cylinders were actuated in the same manner as the wing models and provided the forces due to model weight and inertia that were subtracted from the wing model forces. Details on the robotic apparatus, wing models, dynamic scaling and measurements are available in Supplemental Information, SI7. Force measurements were carried out during four consequent flapping cycles and force data from the third flapping cycle were analyzed to allow the flow to develop and avoid transient phenomena at flapping initiation and termination.

## Acknowledgments

This work was funded by the Israel Science Foundation (ISF), grant No 222/22, to GR

## Data availability

Data used in the study are included in the manuscript and/or supporting information

## List of Symbols and abbreviations

*α*: The mean upstroke inclination angle (Fig. 3) measured on the BRF [°]
*α_p_*: Angle of the upstroke in the XZ plane of the WRF (See Fig. 8E) [°]
*γ*: Climb angle [°]
ϕ: Instantaneous wing sweep angle
ϕ_min_: The upstroke minimum sweep angle (see Fig. 4A)
θ: Instantaneous wing elevation angle [°]
*ζ*: Mean body pitch for the flapping cycle [°]
η: Mean body roll for the flapping cycle [°]
χ: Body angle during flight (see Figs 2 & 9) [°]
*ψ*: Instantaneous wing incidence angle [°]
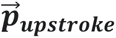: A vector connecting the wingtip positions at upstroke initiation and termination in the BRF
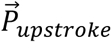: upstroke trajectory vector rotated to the WRF
Re: Reynolds number (based on mean chord and speed of the wing at its radius of gyration
R_X_, R_y_: Matrices for rotation about the X and Y axes, respectively
*Re*: Reynolds number
*t*: Time within the flapping cycle [s]
*T*: Flapping cycle duration [s]
*t*/*T*: No-dimensional time within the flapping cycle
*U_F_*: Instantaneous flight velocity [ms^-1^]
*U̅_F_*: flight velocity averaged across the flapping cycle [ms^-1^]
*U̅*_*F*(*X*)_, *U̅*_*F*(*Y*)_, *U̅*_*F*(*Z*)_: Components of *U̅_F_* along the X, Y and Z axes of the WRF, respectively [ms^-1^]. *U̅_F_*_(*Z*)_is also referred to as the ’climb rate’
*U_W_*: Instantaneous wingtip velocity measured in the WRF [ms^-1^]
*U̅_W_*: cycle averaged mean wingtip velocity
*^U^W*(*Z*): The vertical component of *U_W_* [ms^-1^]
*u_w_*: Instantaneous wingtip velocity relative to the body (measured in the BRF) [ms^-1^]
AoA: Geometrical angle-of-attack of the wing [°]
WRF: World Reference Frame – inertial Cartesian coordinate system (X,Y,Z) where +X is in the horizontal flight direction and +Z is pointing up (opposite gravity)
BRF: Body Reference Frame – non-inertial Cartesian coordinate system (x,y,z) with origin at the thorax where x is the longitudinal axis of the insect, y is the lateral axis and z is the dorsoventral axis (SI4)

## References

Altshuler, D. L., Dickson, W. B., Vance, J. T., Roberts, S. P., & Dickinson, M. H. (2005). Short-amplitude high-frequency wing strokes determine the aerodynamics of honeybee flight. Proceedings of the National Academy of Sciences, 102(50), 18213–18218.

Berg, A. M., & Biewener, A. A. (2008). Kinematics and power requirements of ascending and descending flight in the pigeon (*Columba livia*). Journal of Experimental Biology, 211(7), 1120–1130.

Blackmer, J. L., & Byrne, D. N. (1993). Flight behaviour of *Bemisia tabaci* in a vertical flight chamber: effect of time of day, sex, age and host quality. Physiological Entomology, 18(3), 223–232.

Bramwell, A. R. S., Done, G. & Balmford, D. (2001) Bramwell’s Helicopter Dynamics. Butterworth and Heinemann Publishing. 2^nd^ edition.

Byrne, D. N. (1999). Migration and dispersal by the sweet potato whitefly, *Bemisia tabaci*. Agricultural and Forest Meteorology, 97(4), 309–316.

Cheng, X., & Sun, M. (2018). Very small insects use novel wing flapping and drag principle to generate the weight-supporting vertical force. Journal of Fluid Mechanics, 855, 646–670.

Cheng, X., & Sun, M. (2021). Wing kinematics and aerodynamic forces in miniature insect *Encarsia formosa* in forward flight. Physics of Fluids, 33(2), 021905.

Davidovich, H., & Ribak, G. (2016). Flying with eight wings: inter-sex differences in wingbeat kinematics and aerodynamics during the copulatory flight of damselflies (*Ischnura elegans*). The Science of Nature, 103, 1–8.

Döring, T. F., & Chittka, L. (2007) Visual ecology of aphids a critical review on the role of colours in host finding. Arthropod-Plant Interact, 1, 3–16.

Dudley, R. (2000). The Biomechanics of Insect Flight. Princeton, NJ: Princeton University Press.

Ellington, C. P. (1984a). The aerodynamics of hovering insect flight. III. Kinematics. *Philosophical Transactions of the Royal Society of London. B*, Biological Sciences, 305(1122), 41–78.

Farisenkov, S. E., Kolomenskiy, D., Petrov, P. N., Engels, T., Lapina, N. A., Lehmann, F. O., Onishi, H., Liu, H., & Polilov, A. A. (2022). Novel flight style and light wings boost flight performance of tiny beetles. Nature, 602(7895), 96–100.

Fontaine, E. I., Zabala, F., Dickinson, M. H., & Burdick, J. W. (2009). Wing and body motion during flight initiation in *Drosophila* revealed by automated visual tracking. Journal of Experimental Biology, 212(9), 1307–1323.

Froude, R. E. (1889) On the part played in propulsion by differences of pressure. Transactions of the Institution of Naval Architects Vol 30

Fry, S. N., Sayaman, R., & Dickinson, M. H. (2005). The aerodynamics of hovering flight in *Drosophila*. Journal of Experimental Biology, 208(12), 2303–2318.

Hedrick, T. L. (2008). Software techniques for two- and three-dimensional kinematic measurements of biological and biomimetic systems. Bioinspiration & biomimetics, 3(3), 034001.

Horridge, G. A. (1956). The flight of very small insects. Nature 178, 1334–1335.

Johnson, W. (1980). Helicopter Theory. Princeton University Press. Princeton, New Jersey, USA.

Jones, S. K., Laurenza, R., Hedrick, T. L., Griffith, B. E., & Miller, L. A. (2015). Lift vs. drag based mechanisms for vertical force production in the smallest flying insects. Journal of theoretical biology, 384, 105–120.

KleinHeerenbrink, M., France, L. A., Brighton, C. H., & Taylor, G. K. (2022). Optimization of avian perching manoeuvres. Nature, 607(7917), 91–96.

Krishnan, K., Garde, B., Bennison, A., Cole, N. C., Cole, E. L., Darby, J., … & Shepard, E. L. (2022). The role of wingbeat frequency and amplitude in flight power. Journal of the Royal Society Interface, 19(193), 20220168.

Le Roy, C., Silva, N., Godoy-Diana, R., Debat, V., Llaurens, V., & Muijres, F. T. (2022). Divergence of climbing escape flight performance in Morpho butterflies living in different microhabitats. Journal of Experimental Biology, 225(15), jeb243867.

Lehmann, F.-O., & Dickinson, M.H. (1998). The control of wing kinematics and flight forces in fruit flies (*Drosophila* spp.). Journal of Experimental Biology, 201, 385–401.

Li, Q., Li, H., Shen, H., Yu, Y., He, H., Feng, X., Sun, Y., Mao, Z., Chen, G., Tian, Z., Shen, L., Zheng, X., & Ji, A. (2023). An Aerial–Wall robotic insect that can land, climb, and take off from vertical surfaces. Research, 6, 0144.

Liu, Y., & Sun, M. (2024). Aerodynamics and power requirements of climbing flight in fruit fly model. Physics of Fluids, 36(4).

Lyu, Y. Z., Zhu, H. J., & Sun, M. (2019). Flapping-mode changes and aerodynamic mechanisms in miniature insects. Physical review E, 99(1), 012419.

MacAyeal, L. C., Riskin, D. K., Swartz, S. M., & Breuer, K. S. (2011). Climbing flight performance and load carrying in lesser dog-faced fruit bats (*Cynopterus brachyotis*). Journal of Experimental Biology, 214(5), 786–793.

Marden, J. H. (1987). Maximum lift production during takeoff in flying animals. Journal of experimental Biology, 130(1), 235–258.

Meng, X., Liu, Y., & Sun, M. (2017). Aerodynamics of ascending flight in fruit flies. Journal of Bionic Engineering, 14(1), 75–87.

Nguyen, A. T., Le, V. D. T., Tran, T. H., Duc, V. N., & Phung, V. B. (2020). Study of vertically ascending flight of a hawkmoth model. Acta Mechanica Sinica, 36, 1031–1045.

Ortega-Jiménez, V. M., & Dudley, R. (2018). Ascending flight and decelerating vertical glides in Anna’s hummingbirds. Journal of Experimental Biology, 221(24), jeb191171.

Parslew, B. (2015). Predicting power-optimal kinematics of avian wings. Journal of The Royal Society Interface, 12(102), 20140953.

Perring, T. M., Stansly, P. A., Liu, T. X., Smith, H. A., & Andreason, S. A. (2018). Whiteflies: Biology, Ecology, and Management. Sustainable management of arthropod pests of tomato, pp. 73–110. Academic Press.

Rayner, J. M. V., & Aldridge, H. D. J. N. (1985). Three-dimensional reconstruction of animal flight paths and the turning flight of microchiropteran bats. Journal of Experimental Biology, 118, 247–265.

Ribak, G., Dafni, E., & Gerling, D. (2016). Whiteflies stabilize their take-off with closed wings. Journal of Experimental Biology, 219(11), 1639–1648.

Sane, S. P. (2016). Neurobiology and biomechanics of flight in miniature insects. Current opinion in neurobiology, 41, 158–166.

Sarig, A., & Ribak, G. (2021). To what extent can the tiny parasitoid wasps, *Eretmocerus mundus*, fly upwind?. Journal Applied Entomology. 2021(00), 1– 15.

Sun, M. (2023). Colloquium: Miniature insect flight. Reviews of Modern Physics, 95(4), 041001.

Vogel, S. (1988). In: Life’s Devices: The physical world of animals and plants (pp. 152–154): Princeton University Press.

Wainwright, C. E., Reynolds, D. R., & Reynolds, A. M. (2020). Linking small-scale flight manoeuvres and density profiles to the vertical movement of insects in the nocturnal stable boundary layer. Scientific reports, 10(1), 1019.

Wainwright, C. E., Stepanian, P. M., Reynolds, D. R., & Reynolds, A. M. (2017). The movement of small insects in the convective boundary layer: linking patterns to processes. Scientific reports, 7(1), 5438.

Weis-Fogh, T. (1973). Quick estimates of flight fitness in hovering animals, including novel mechanisms for lift production. Journal of experimental Biology, 59(1), 169–230.

Williams, E. J. Murray-Bruce, J. & Murphy, D. M. (2025) An inverse problems approach to micro-PIV for measuring flow around freely flying tiny insects. Journal of Experimental Biology 228:jeb249417

