## Supporting information (text, figures and tables)) for "Backstroke rowing as an adaptation for low-Reynolds-number flight in tiny whiteflies (*Bemisia tabaci*)"

**Content:**

- Supplemental Movie Caption
- SI1 – Supplementary Table S1.1 and Table S1.2. Flight parameters of the analyzed flapping cycles and their correlation with flight velocity.
- SI2 - Prevalence of upside-down flight postures during climb
- SI3 - Wingtip trajectories in the world reference frame (WRF) for the 15 movies not included in Fig. 8 of the main text
- SI4 - Observed *vs.* calculated (eqs 1 &2 in the main text) inclination of the upstroke in the WRF
- SI5 - Relationship between forewing and hindwing kinematics
- SI6 - Extraction of flapping kinematics
- SI7 - Robotic force measurements

### Supplemental Movie caption:

**Movie 1 - Example of the high-speed films analyzed.** The movie shows an example of flight captured by three spatially-calibrated high-speed cameras. The insect in the film is ascending at a climb angle of 60° and a climb rate of 0.161 m/s.

### SI1 - Flight parameters of the analyzed flapping cycles (Table S1.1) and their correlation with flight velocity (Table S1.2)

**Table S1.1 Data means for the analyzed flapping cycles.** $\overline{U}_{F(X)}$ = horizontal body speed. $\overline{U}_{F(Z)}$= vertical body speed, $\overline{U}_{F}$ = three-dimensional body speed, $\gamma$ = climb angle, χ = flight body angle. η= body roll, f = wingbeat frequency J = advance-ratio, *Re* = Reynolds number calculated as $\bar{c}\overline{U_{rg}}/\upsilon^{-1}$, where $\overline{U_{rg}}$ is the mean wing speed measured at the radius of gyration (0.853R), $\bar{c}$ the mean wing chord (0.26R) and υ the kinematic viscosity of air ($15 \times 10^{-6}$ $m^2s^{-1}$).

| Climb group | $\overline{U}_{F(X)}$ ($ms^{-1}$) | $\overline{U}_{F(Z)}$ ($ms^{-1}$) | $\overline{U}_{F}$ ($ms^{-1}$) | γ (°) | χ (°) | η (°) | f (Hz) | J | Re |
|---|---|---|---|---|---|---|---|---|---|
| **30°** | 0.367 | 0.215 | 0.426 | 30 | 40 | 57 | 214 | 0.41 | 14 |
| | 0.193 | 0.098 | 0.217 | 27 | 55 | 84 | 182 | 0.22 | 15 |
| | 0.308 | 0.166 | 0.350 | 28 | 150 | -124 | 194 | 0.26 | 17 |
| | 0.401 | 0.261 | 0.479 | 33 | 43 | -93 | 194 | 0.37 | 21 |
| | 0.179 | 0.088 | 0.199 | 26 | 62 | 85 | 176 | 0.17 | 18 |
| | 0.228 | 0.138 | 0.267 | 31 | 125 | -115 | 200 | 0.26 | 17 |
| **45°** | 0.194 | 0.193 | 0.273 | 45 | 61 | 75 | 188 | 0.21 | 23 |
| | 0.228 | 0.204 | 0.305 | 42 | 65 | 55 | 194 | 0.23 | 20 |
| | 0.314 | 0.348 | 0.469 | 48 | 141 | -148 | 207 | 0.37 | 17 |
| | 0.299 | 0.263 | 0.399 | 41 | 55 | 58 | 194 | 0.36 | 18 |
| | 0.164 | 0.140 | 0.216 | 40 | 134 | 127 | 214 | 0.21 | 13 |
| | 0.137 | 0.122 | 0.184 | 42 | 152 | -137 | 214 | 0.18 | 13 |
| **60°** | 0.156 | 0.251 | 0.295 | 58 | 104 | 56 | 194 | 0.23 | 20 |
| | 0.150 | 0.237 | 0.281 | 58 | 109 | -17 | 188 | 0.19 | 23 |
| | 0.056 | 0.144 | 0.155 | 69 | 137 | 158 | 207 | 0.14 | 14 |
| | 0.263 | 0.419 | 0.494 | 58 | 137 | 153 | 207 | 0.36 | 19 |
| | 0.180 | 0.300 | 0.350 | 59 | 122 | -116 | 167 | 0.28 | 20 |
| | 0.094 | 0.161 | 0.187 | 60 | 135 | 11 | 207 | 0.15 | 20 |

**Table S1.2 correlation matrix between climb performance and flapping kinematics parameters**

Pearson correlations are presented with significant correlations denoted in bold red font and asterisks denote significance level (* p<0.05, ** p<0.01). $\overline{U}_F$, $\overline{U}_{F(X)}$, $\overline{U}_{F(Z)}$ and $\gamma$ are the 3D flight speed, horizontal flight speed, climb rate and climb angle, respectively. They are tested for correlation with: flapping frequency (*f*), body flight angle ($\chi$), body roll ($\eta$), body pitch ($\zeta$), downstroke minimum in sweep angle (at $t/T$ ≈0.25, $\phi_d$), sweep angle minimum during the upstroke ($\phi_{min}$). $\theta_{min}$ and $\theta_{max}$ are the minimum and maximum values of the elevation angle, $\psi_d$ and $\psi_{up}$ are the incidence angle during the downstroke and upstroke, respectively, AoA is the angle of attack during $t/T$=0.6 of the upstroke, $u_w$ is the speed of the wingtip relative to the body at the same instance and $U_{W(Z)}$ is the vertical component of the wingtip velocity in the WRF at the same instance. $\alpha$ and $\alpha_p$ are the inclination of the upstroke in the BRF and WRF, respectively. $Acc_{W(Z)}$ is the vertical component of wing acceleration during the upstroke in the WRF

| | $f$ | $\chi$ | $\eta$ | $\zeta$ | $\phi_d$ | $\phi_{min}$ | $\theta_{min}$ | $\theta_{max}$ | $\psi_d$ | $\psi_{up}$ | $AoA$ |
|---|---|---|---|---|---|---|---|---|---|---|---|
| $\overline{U}_F$ | 0.04 | -0.22 | 0.14 | -0.16 | -0.03 | **-0.57*** | -0.34 | -0.25 | 0.02 | -0.11 | -0.16 |
| $\overline{U}_{F(X)}$ | 0.05 | -0.43 | 0.02 | -0.28 | 0.22 | -0.47 | -0.21 | -0.34 | 0.11 | -0.09 | -0.16 |
| $\overline{U}_{F(Z)}$ | 0.02 | 0.09 | 0.15 | 0.02 | -0.27 | **-0.48*** | -0.37 | -0.08 | -0.11 | -0.06 | -0.12 |
| $\gamma$ | 0.1 | **0.47*** | 0.01 | 0.19 | -0.44 | -0.05 | 0.01 | 0.29 | -0.18 | 0.11 | 0.18 |

| | $u_w$ | $U_{W(Z)}$ | $\alpha$ | $\alpha_p$ | $Acc_{W(Z)}$ |
|---|---|---|---|---|---|
| $\overline{U}_F$ | 0.43 | 0.05 | 0.44 | 0.15 | -0.11 |
| $\overline{U}_{F(X)}$ | 0.18 | 0.36 | 0.31 | 0.45 | 0.19 |
| $\overline{U}_{F(Z)}$ | **0.61**** | -0.33 | 0.43 | -0.25 | -0.40 |
| $\gamma$ | 0.33 | **-0.65** | 0.17 | **-0.69**** | **-0.56*** |

### SI2 Prevalence of upside-down flight postures during climb

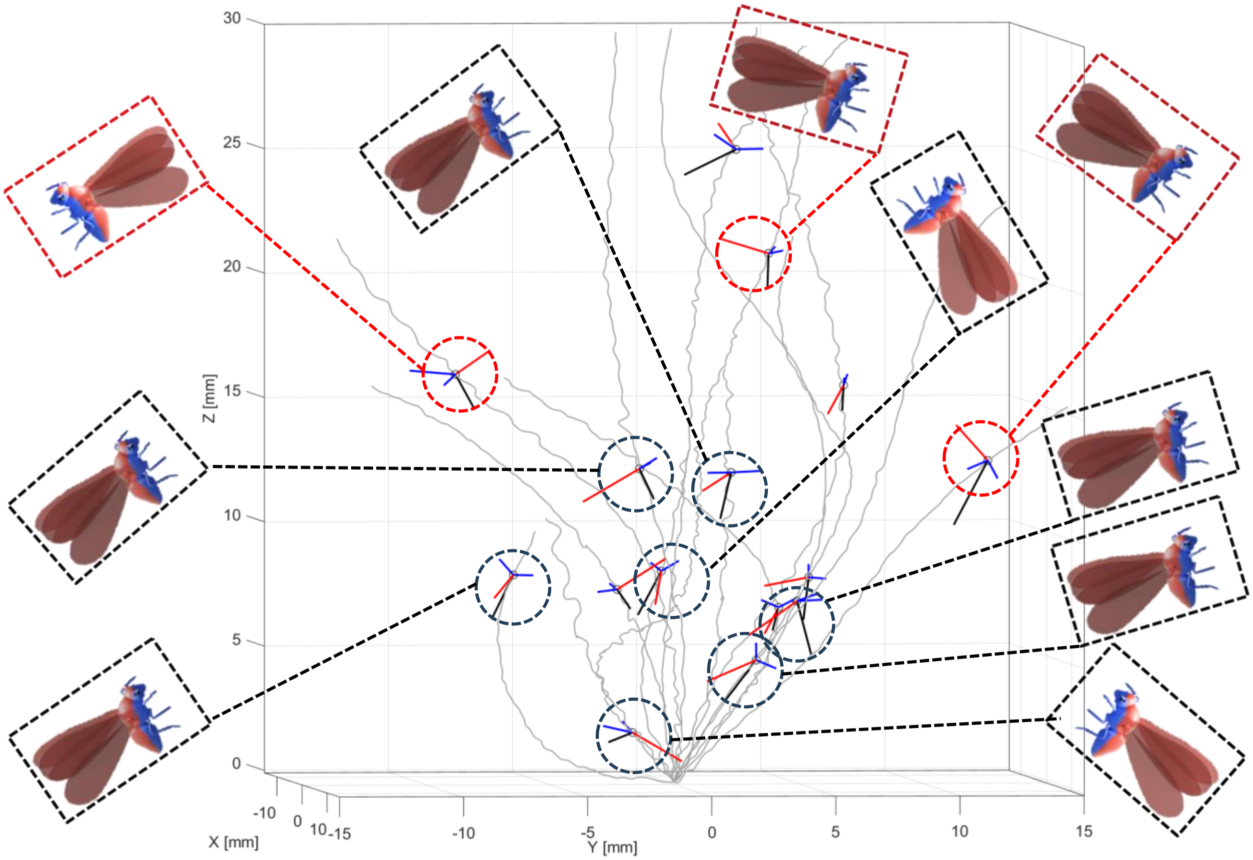


**Figure SI 1. Occurrence of upside-down body posture during climb.** Fifteen flight trajectories at different climb angles are shown in grey. The orientation of the insects is illustrated with lines denoting the thorax longitudinal axis (black), the wings in the dorsal stroke reversal (during the clap, red), and the wings in the ventral stroke reversal (blue). For some of the insects the 3D orientation is illustrated in 2d using animated insects where red dashed frame lines show insects in the right-side up orientation (the dorsal side is pointing up and the body is in an acute angle), whereas dashed black frames denote upside-down (the wingtips point down during the clap and are beneath the wingtips during the ventral stroke reversal).

**SI3 - Wingtip trajectories in the world reference frame (WRF) for the 15 movies not included in Fig. 8 of the main text**

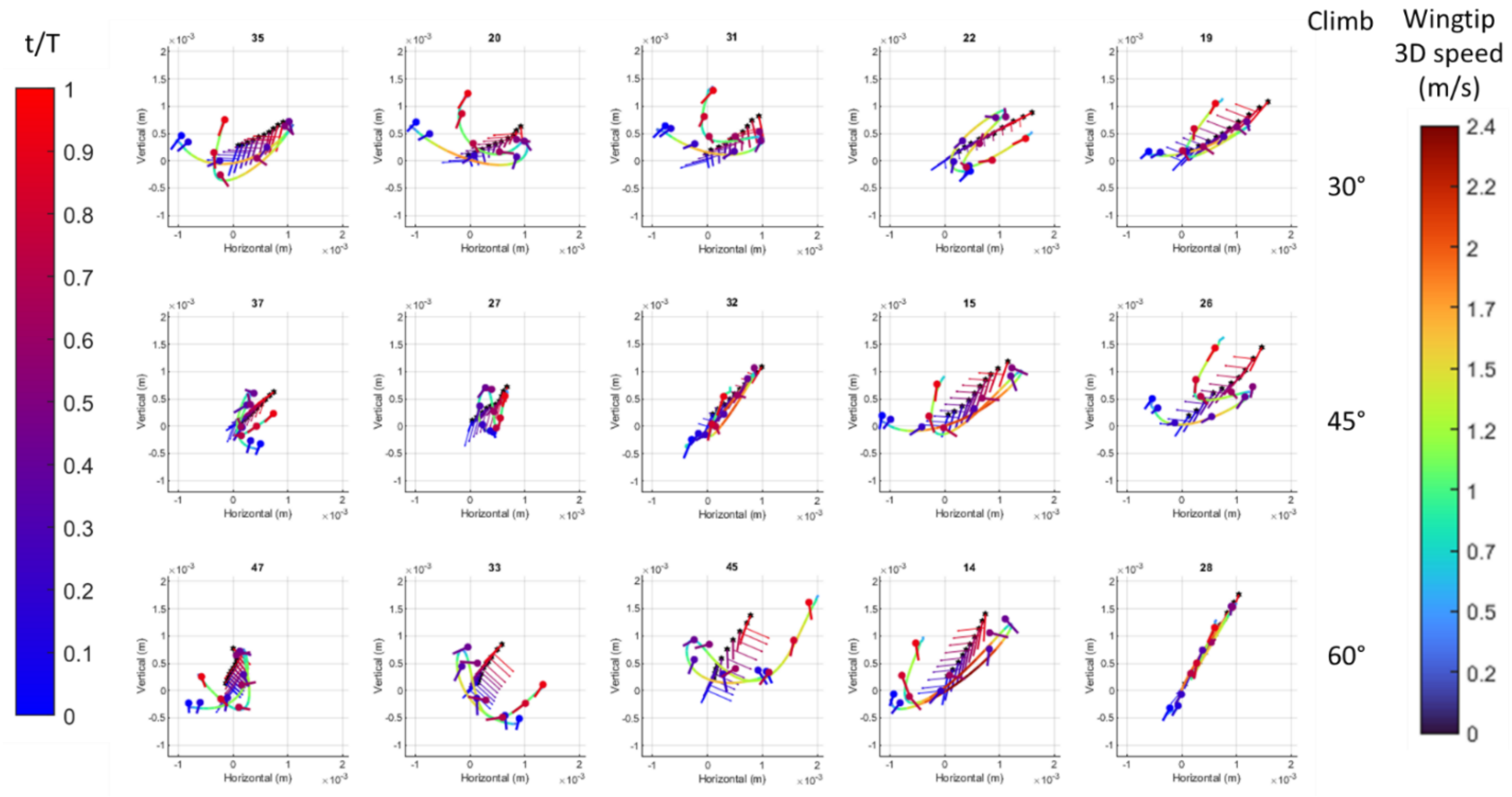

**Figure SI 2 (previous page): Wingtip trajectories in the world reference frame**. All analyzed flapping cycles are shown, except the three presented in Fig. 8 of the main text. The cycles are arranged according to climb angle category (rows) and increasing 3D flight speed (left to right). The data are rotated with the positive side of the horizontal axis pointing in the horizontal flight direction Wingtip trajectory is color-coded according to wingtip three-dimensional speed (right color-bar). In each plot the body and wing chord are shown at 0.1 t/T interval and color-coded according to non-dimensional time within the flapping cycle (t/T, left color-bar). Chords are denoted by thick lines with circles denoting the leading edge. Thinner lines denote the longitudinal body axis, with black hexagons denoting the head and arrows indicating the insect's dorsal side. For a high-resolution presentation see link.

#### SI4 - Observed *vs.* calculated (eqs 1 &2 in the main text) inclination of the upstroke in the WRF

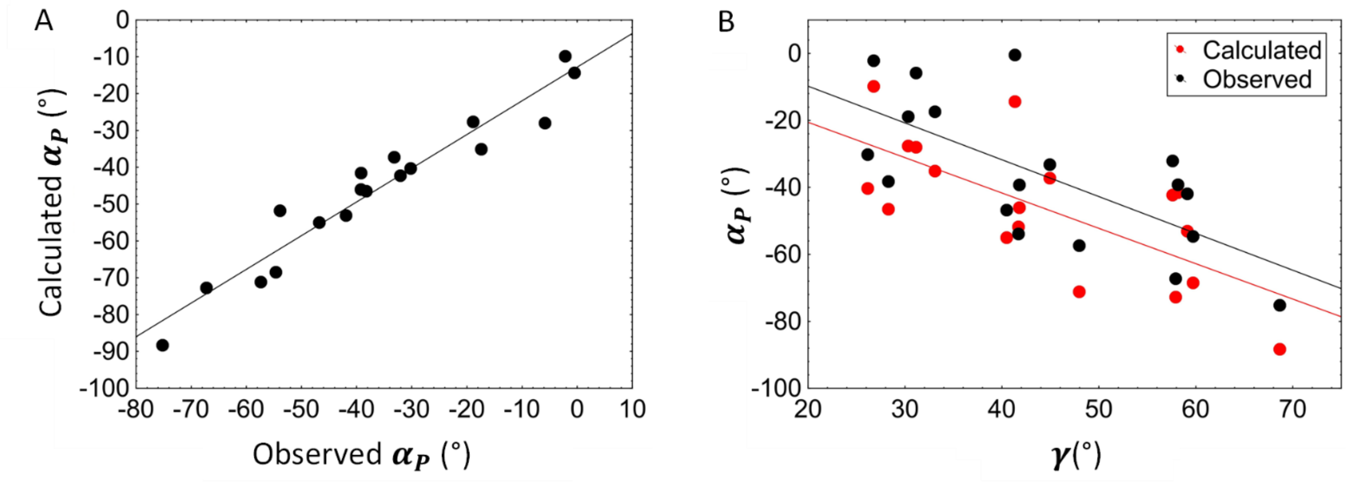


**Figure SI 3: Comparison between observed and calculated upstroke inclination in the WRF.** A) Correlation between the observed upstroke direction (**$\alpha_P$**) in the films and the upstroke direction calculated from the body roll and pitch (eq. 2, 3 in the main text), illustrating that a combination of all three parameters in eq. 2 are required to explain ($R^2 = 0.93$) the observed direction (correlation observed versus predicted: r=0.965, p<0.0001). B) both the observed and calculated $\alpha_P$ are negatively correlated with climb angle ($\gamma$).

#### SI5 - Relationship between forewing and hindwing kinematics

In the movies the hindwings lag behind the forewings with a phase shift of ~0.1$T$ (were $T$ is the flapping cycle duration) and an offset of ~+20° in the sweep angle and ~-20° in the elevation angle. In addition, the incidence angle of the hindwings reaches a lower minimum and a higher maximum relative to the forewings. Since these differences between the ipsilateral wings remained consistent across different films (Fig. SI 4) the main text focuses on describing only the motion of the forewing.

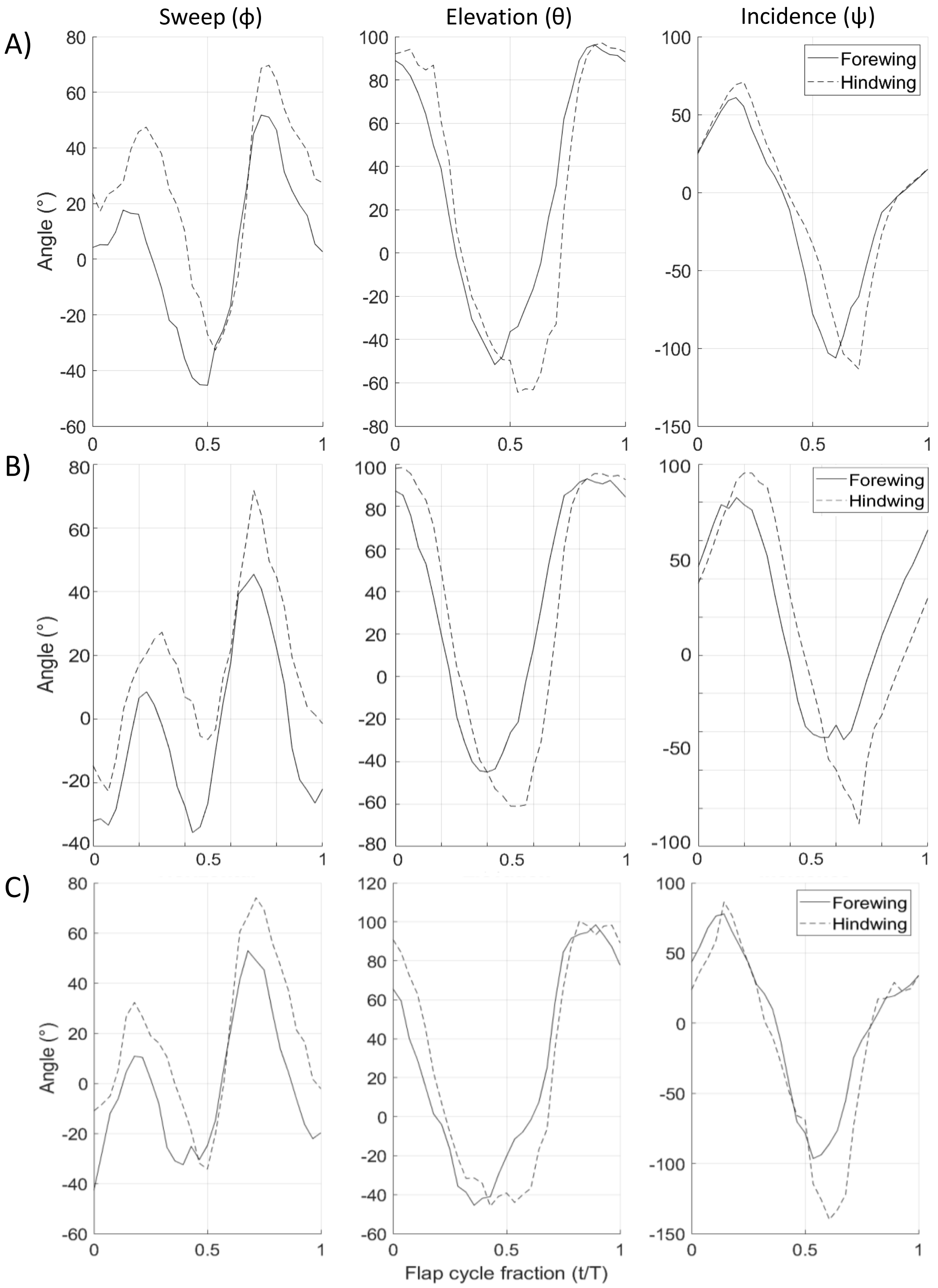


**Figure SI 4: Forewing and hindwing kinematics.** Columns display the three Euler angles of the fore- and hind-wing in three of the analyzed sequences (rows). The horizontal axis in non-dimensional time within the flapping cycle. The motion of the hindwings (dashed lines) principally tracks the motion of the forewing (solid lines) with a phase different of ~0.1T.

**SI6 - Extraction of flapping kinematics**

*Transformation between WRF and BRF*

The extraction of flapping kinematics begins with transformation of the data from the camera frame of reference to the body reference frame (BRF) using the following steps:

1) In the first video frame, the center of the thorax in the camera frame of reference is defined as the origin, and the coordinate system is rotated about the Z axis to align +X with the horizontal flight direction. This coordinate system is hereafter referred to as the world reference frame (WRF).

2) In each video frame, the origin of the WRF is shifted to the center of the insect's thorax. Then, using the points on head, posterior tip of the thorax and the two wing hinges, we found the direction cosines of the anatomical body axes in the WRF. If $X_B$ and $Y_B$ are unit vectors in the direction of the longitudinal and transverse body axes (Fig. SI 2A) in the WRF, respectively, then $Z_B$ is defined as the dorsoventral axis by the cross-product of $X_B$ and $Y_B$:

$$\widehat{Z_B} = \frac{\widehat{X_B} \times \widehat{Y_B}}{sin\ \epsilon}, \qquad \text{(SI eq. 1)}$$

in which the cap above the vectors denotes a unit vector. $\epsilon$ is the angle between $X_B$ and $Y_B$, which should be 90° but might slightly vary due to digitization errors. We ensured that all resulting axes were orthogonal by recalculating $Y_B$ based on $\widehat{Z_B}$ and $\widehat{X_B}$:

$$\widehat{Y_B} = \widehat{Z_B} \times \widehat{X_B}. \qquad \text{(SI eq. 2)}$$

3) The unit vectors of the body axes are then used to form the rotation matrix M, written as:

$$\mathbf{M} = \begin{bmatrix} \widehat{X_B} \\ \widehat{Y_B} \\ \widehat{Z_B} \end{bmatrix} \qquad \text{(SI eq. 3)}$$

The matrix rotates the position ($P$) of each point on the body and wing from the (WRF) to the body frame of reference (BRF):

$$P_B = M \times P\ , \qquad \text{(SI eq. 4)}$$

with $P_B$ denoting the position of P in the BRF.

*Reconstruction of wing flapping angles*

Wing-flapping angles were calculated in the BRF as follows:

The origin is shifted to the position of the relevant wing hinge. The elevation angle of the wing (θ) was calculated for each frame in the body's frontal (yz, Fig. SI 5A) plane as:

$$\theta = \tan^{-1}\left(\frac{WL_z}{WL_y}\right) \qquad \text{(SI eq. 5)}$$

for the left wing and

$$\theta = \tan^{-1}\left(\frac{WL_z}{-WL_y}\right) \qquad \text{(SI eq. 6)}$$

for the right wing, with $WL$ being the wings' length axis (Fig. SI 5B) and the subscripts y and z denoting the relevant components in the body frame of reference. The sweep angle ($\phi$) in the horizontal axis was calculated for each frame by rotating the wing position data ($P_1$) about the longitudinal axis by $\theta$, so that the wing axis lies is in the body plane (xy):

$$P_1' = \begin{bmatrix} 1 & 0 & 0 \\ 0 & \cos\theta & \sin\theta \\ 0 & -\sin\theta & \cos\theta \end{bmatrix} \times P_1\ . \qquad \text{(SI eq. 7)}$$

Sweep angle ($\phi$) was calculated from the rotated data of the left and right wings respectively as:

$$\phi = \tan^{-1}\left(\frac{WL_x}{WL_y}\right) \text{ and } \phi = \tan^{-1}\left(\frac{WL_x}{-WL_y}\right) \qquad \text{(SI eqs 8 \& 9)}$$

The angle of incidence ($\psi$) was calculated following an additional rotation of the wing data about the insect's dorsoventral axis by $\phi$ to align the wing axis with the y-axis. This was achieved using:

$$P_1'' = \begin{bmatrix} \cos\phi & -\sin\phi & 0 \\ \sin\phi & \cos\phi & 0 \\ 0 & 0 & 1 \end{bmatrix} \times P_1'. \qquad \text{(SI eq. 10)}$$

The angle of incidence is thus:

$$\psi = -\tan^{-1}\left(\frac{WC_z}{WC_x}\right). \qquad \text{(SI eq. 11)}$$

The wing's angle-of-attack (AoA) with respect to the direction of wing motion is calculated from wing length ($WL$) and chord ($WC$) vectors as illustrated in Fig. SI 5B. LV and FV are vectors connecting the wing hinge with leading and trailing edge at 0.7 of the wing length, respectively. The vector (WN) normal to the area of the left wing is calculated from the following cross-product:

$$WN = LV \times TV \ . \qquad \text{(SI eq. 12)}$$

The vector (WSN) perpendicular to both wing velocity (WV) and the wing length (WL) is:

$$WSN = WV \times WL \qquad \text{(SI eq. 13)}$$

and AoA is derived as the angle between $WN$ and $WSN$ from their dot product:

$$AoA = \cos^{-1}\left(\frac{WN \cdot WSN}{|WN| \cdot |WSN|}\right). \qquad \text{(SI eq. 14)}$$

Calculations for the right wing were similar but the order of the cross-products was changed according to the right-hand rule.

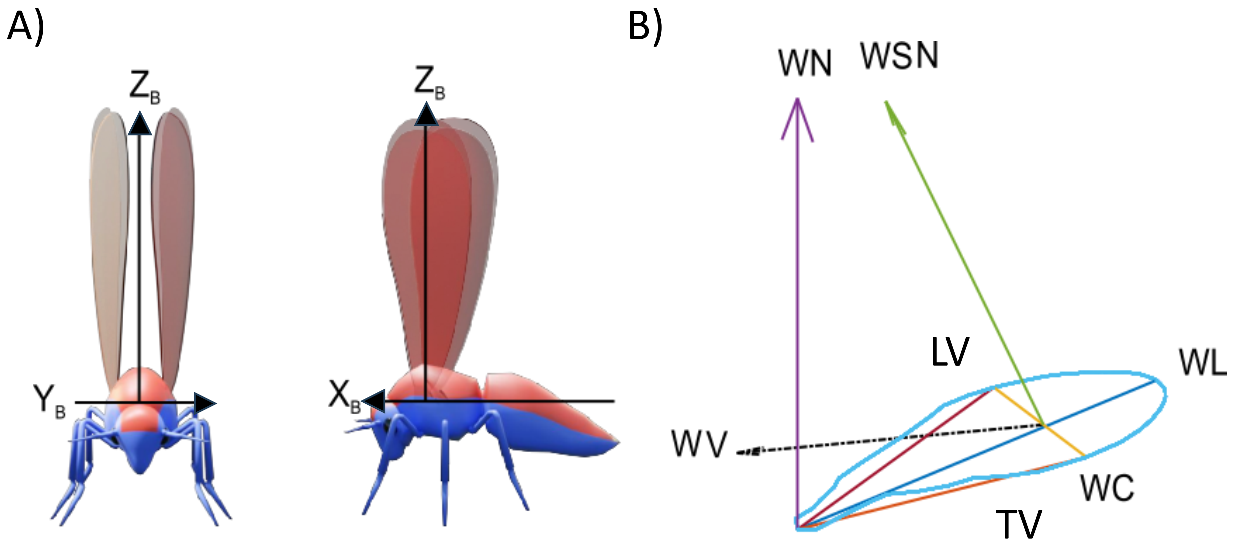


**Figure SI 5: Definitions and reconstruction of body and wing kinematics.** A) Body axes used to rotate the coordinate system from the WRF to the BRF. B) Coordinate system and positions on the wing used to find the angle-of-attack. WL is wing length between tip and hinge, WC is wing chord between leading and trailing edges at 0.7 of wing length, and $Z_W$ is a cross-product of vectors indicating the leading (LV) and trailing (TV) edges. WN is the vector normal to the wing area (pointing towards the wing's dorsal side.

WV is the wing velocity vector (shown here as a dashed black arrow). WSN (green) is a vector normal to both wing length and wing velocity.

### SI 7 Robotic force measurements

To determine the contribution of the upstroke rowing motion to upthrust generation we replicated the flapping motion on a 290:1 upscaled whitefly wing model connected at the wing root to a tri-axes force transducer (K3D60, Me-systems). The other end of the force transducer was connected to the tip of a custom-built robotic system that actuated two wing models through Glycerin.

*Wing model* – The wing models are based on ×290 magnification of the whitefly forewing planform (Figure 1C in Ribak et al. 2016) converted to a 3D CAD model in Solidworks®. The model included spanwise and chordwise gradients of diminishing thickness with a minimal thickness of 4 mm. The thickness at the wing base was 21 mm to fit the force transducer. From this base the thickness diminished spanwise with a curvature of 0.424 $m^{-1}$. Consequently, constant minimal wing thickness was reached at 0.69 of the wing length and remained constant further towards the tip (Fig SI 6). The chordwise thickness gradient ensured the proximal trailing edge had a lower thickness than the leading edge. At 19 mm from the base (6.6% of the wing length) the leading and trailing edges had thicknesses of 15.2 and 12.6 mm respectively. At 33.3% from the base the thickness was 8.4 and 6.6 mm in the leading and trailing edges respectively. These thickness gradients were inspired by the mass distribution and thickness gradients present in real insect wings but they were modified to provide a smooth continuous surface with minimal thickness at the trailing edge while preventing bending and breaking of the model under the load of the fluid force. The base of the wings fitted the force transducer so that the axis of wing flapping concurred with the position of the insect's wing hinge.

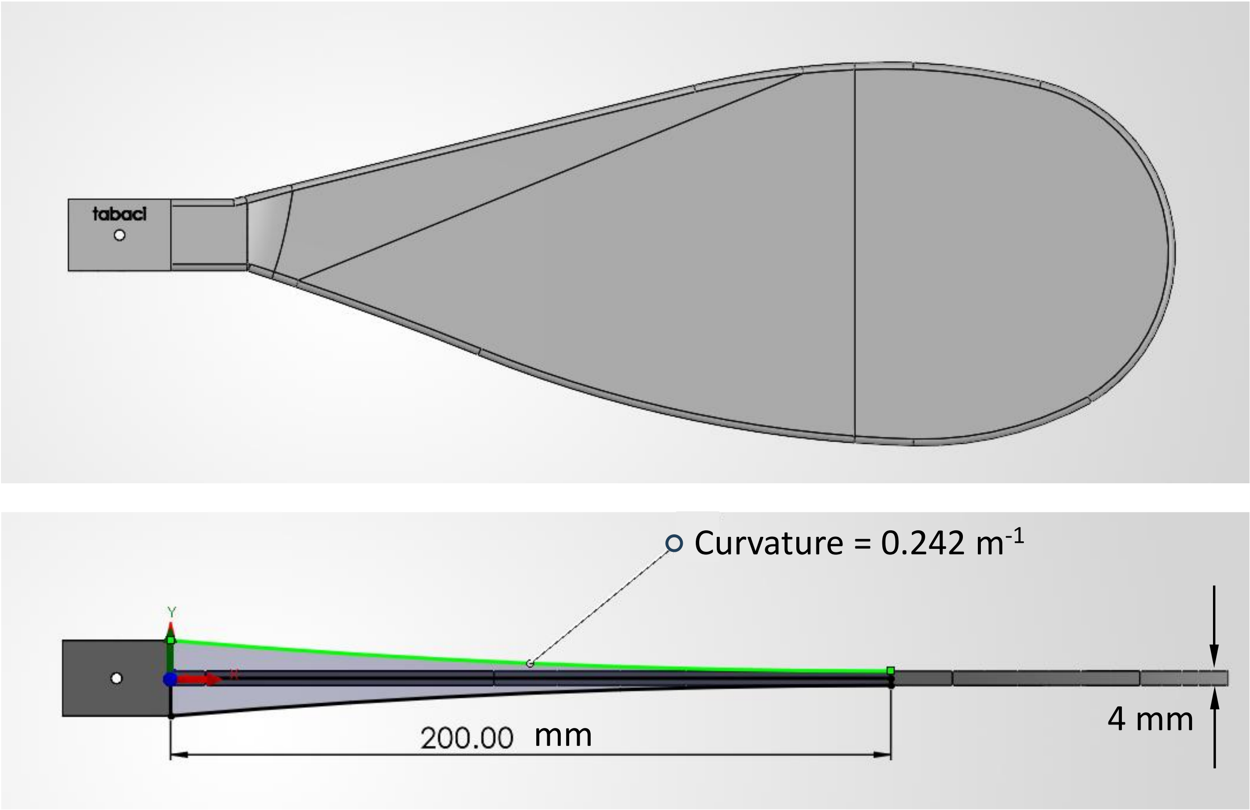


**Figure SI 6 CAD sketches of the wing model**. shown are the rectangular base fixed to the force transducer, and the planform (upper panel) and thickness gradients at the leading edge (lower panel). The spanwise thickness gradient ends 20 cm from the wing base and the remaining wing has a constant thickness of 4 mm. Similarly, the thickness in the proximal 0.69 of the wing length declines from the leading to the trailing edge.

*Robotic system* - The robotic set-up included two Nema 23 step motors (M1262E106, LAM Technologies, Italy) and a set of pulleys and gears (Fig SI 7) that flapped the two wing models in bilaterally symmetric fashion. Two closed loop drivers (DDS1, LAM Technologies, Italy) and a programable controller (PMX-4EX-SA, Arcus Technology, USA) allowed to dictate the motion of the wing models simultaneously in two degrees-of-freedom: the flapping ($\phi$) angle in a horizontal stroke plane and rotation of the wing model about its span (angle-of-attack). The controllers controlled both the position and angular speed of both angles at each instant. The wing models were submerged 0.4 m below the surface in a 1.0 × 1.0 m, 0.8 m deep glycerin tank. The data from the force transducer was recorded on a desktop computer at 100 measurements per second.

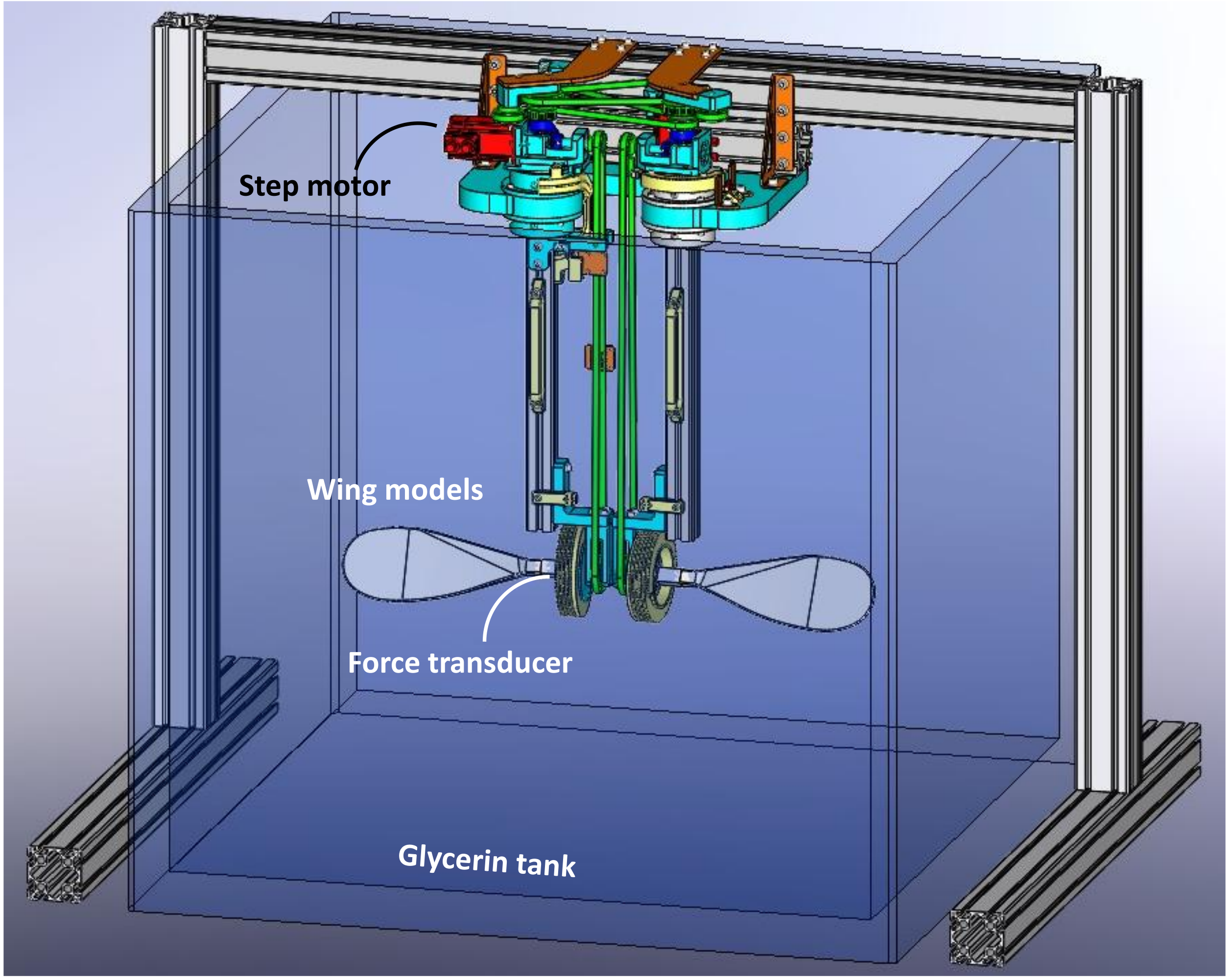


**Figure SI 7 The robotic setup with the wing models.** The wings flap about the vertical axis passing through a position on the wing that is equivalent to the anatomical hinge in the insects.

*Robotic wings kinematics* – The 3D flapping kinematics of the real insects was converted into robotic actuation with 2 degrees-of-freedom by "flattening" the 3D wing tip trajectory to a horizontal stroke plane while conserving the magnitude of the 3D wing speed and angle of attack. This was achieved by converting the three instantaneous Euler angles (SI2) into instantaneous wing velocity and angle of attack. The process then applies a piece-wise integration of the velocity to determine the instantaneous flapping angle in the flat flapping plane, at each instance. Concurrently, the instantaneous AoA of the wing model is due to a combination of the instantaneous wing incidence and elevation angles. An example of the resulting flattened robot kinematics for the case of 30° climb angle is shown in Fig. 9B of the main text. The kinematics is converted to a program for the robot to flap the wings for four consecutive flapping cycles. Three such programs were written, each corresponding to the mean flapping kinematics of the insects in the 30°, 45° and 60° climb angle.

*Dynamic Similarity* – for the force measurements to represent flow conditions similar to miniature insects, the motion of the wing models was dynamically scaled to have the same Re numbers as the insects. The kinematic viscosity of the Glycerin in the tank was determined using a Cannon-Fenske glass viscometer. The force measurements were performed in a climate-controlled room (25° ± 1°) to avoid temperature fluctuations from changing the viscosity of the Glycerin during the measurements. The speed of the wing models was adjusted to match the Re numbers of the insect wing according to:

$$\frac{U_{robot}}{U_{insect}} = \frac{\nu_{glycerine}}{\nu_{air}} \frac{R_{insect}}{R_{robot}} \quad \text{(SI eq. 15)}$$

Where $\nu_{gly}$ is the kinematic viscosity of the Glycerin in the experiment ($\nu_{gly}$ = 4.5×10$^{-4}$ m$^2$s$^{-1}$), $\nu_{air}$ is the kinematic viscosity of air (15×10$^{-6}$ m$^2$s$^{-1}$), $R_{insect}$ and $R_{robot}$ are the wing lengths of the insect and robot, respectively and $U_{robot}$ and $U_{insect}$ are the speed of the robot and insect wing.

*Analysis of force data* – After performing four consecutive flapping cycles at the mean kinematics of the insects from the 30°, 45°, and 60° climb angle categories, the measurements were repeated with cylindrical models replacing the wing models. These cylinders had the same mass, density and distance of the center of mass from the axis of flapping as the wing models. The force data from the cylinders were used to correct for model buoyancy and inertia in the wing force measurements as follows:

The forces from the force transducer were converted to vertical and horizontal forces using the known angle of attack ($\alpha$). The drag force ($D$) was obtained from the horizontal force ($F_H$). At any instant:

$$D = F_H - (F_i + F_a \cos\alpha) \quad \text{(SI eq. 16)}$$

where $F_i$ is the inertia of the wing model found from blade-element analysis:

$$F_i = \ddot{\phi} \int_0^R r m_{(r)} dr \quad \text{(SI eq. 17)}$$

$\ddot{\phi}$ is the instantaneous angular acceleration of the wing model due to the flapping motion. $F_a$ is the added mass of the wing model calculated as in Sane et al. (2002).

$$F_a = \frac{\rho \pi R}{4} \left[ \left( \ddot{\phi} \sin\alpha + \dot{\phi}\dot{\alpha} \cos\alpha \right) \int_0^R r c_{(r)}^2 dr - \frac{\ddot{\alpha}\bar{c}}{4} \int_0^R c_{(r)}^2 \, dr) \right] \quad \text{(SI eq. 18)}$$

$F_a$ is normal to the wing chord. Therefore, it is divided into its horizontal and vertical component using the angle-of-attack.

The lift force ($L$) is calculated from the vertical force ($F_V$) by subtracting the measured vertical cylinder force from the measured vertical wing model. This operation accounts for the model weight in glycerin. From the remaining force we subtract the vertical component of $F_a$:

$$L = F_{V(wing)} - (F_{V(cylinder)} + F_a \sin\alpha) \quad \text{(SI eq. 19)}$$

To find the contribution of lift and drag to supporting the body weight in air during the upstroke rowing phase, we focus on the forces from the beginning of the third upstroke to the point where the wing reaches the sweep minimum $\phi_{min}$ (2.5≤ $t/T$ ≤2.75) in the third flapping cycle. The focus on the third flapping cycle is a precaution to allow the flow to develop and prevent transient phenomena at the start and end of the experiment. Fig 9C in the main text shows, however, that the measurements are quite repeatable even from the first flapping cycle.

The contribution of lift and drag to vertical force in the WRF is:

$$F_{(Z)} = D \sin\alpha_p + L \cos\alpha_p \quad \text{(SI eq. 20)}$$

where $\alpha_p$ from the movies is 18.8°, 38.5° and 51.7° degrees in the 30°, 45° and 60° climb angle categories, respectively. The instantaneous forces during the upstroke rowing (2.5≤ $t/T$ ≤2.75) are averaged. To scale the measured forces back to the forces encountered by the insect wing:

$$F_{inset} = 4F_{robot} \frac{\left(\frac{R_{robot}}{R_{insect}} \frac{\nu_{air}}{\nu_{glycerine}}\right)^2}{\frac{\rho_{robot}}{\rho_{insect}} \left(\frac{R_{robot}}{R_{insect}}\right)^2} \quad \text{(SI eq. 21)}$$

where the multiplication by 4 is added to account for the four wings of whiteflies. The numerator is $\left(\frac{U_{insect}}{U_{robot}}\right)^2$ and the denominator corrects for differences in density ($\rho$) between media (glycerin versus air) and the larger area of the robotic wing model. These vertical forces of the insect are the ones presented for the three different climb angles as stacked bars in Fig 9E of the main text.
